# Short Term Evolutionary Dynamics of *Escherichia Coli* in Different Carbon Environments

**DOI:** 10.1101/398578

**Authors:** Debika Choudhury, Supreet Saini

**Affiliations:** Department of Chemical Engineering, Indian Institute of Technology Bombay, Mumbai, 400076, India

**Keywords:** Sugars, *E. coli*, evolution, metabolism

## Abstract

Starting from a parental*E. coli* K-12 MG1655 strain, we evolve cells in five different carbon environments-glucose, arabinose, xylose, rhamnose, and a mixture of these four sugars (in a predefined ratio) for approximately 2,000 generations. At the end of the adaptation period, we quantify and compare growth dynamics of the strains in a variety of environments. The evolved strains show no specialized adaptation towards growth in the carbon medium in which they were evolved. Rather, in all environments, the evolved strains exhibited a reduced lag phase and an increased growth rate. Sequencing results reveal that these dynamical properties are not introduced via mutations in the precise loci associated with utilization of the sugar in which the bacterium was evolved in. These phenotypic changes are rather likely introduced via mutationselsewhere onthe genome. Sugar systems are known to exhibit hierarchy in utilization. Evolution in a defined environment, in our experimental framework, does not alter this hierarchy.

## 1. Introduction

Bacteria are capable of utilizing a wide range of sugar molecules as sources of carbon and energy(Mayer and Boos 2005). Utilization systems specific for each of these sugars exist and consist of a set of transporters which mediate uptake of the specific sugar from the external medium, a set of enzymes which convert the internalized sugar into metabolic intermediate and a regulator which controls the overall expression of the transporter and metabolic genes. The latter genes are not always expressed; rather subjected to conditional induction. In presence of cognate sugar (inducer), the regulatory protein activates transcription of the utilization genes. When the inducer is absent, the same protein acts as a negative regulator, turning off expression of the transporter and metabolic genes(Afroz, Biliouris et al. 2014).Thus this regulatory strategy is a cost-saving one, whereby cells invest in synthesis of utilization machinery only when needed rather than allowing unnecessary production (Dekel and Alon 2005).Global regulators, such as CRP, also control expression of the sugar utilization genes(Shimada, Fujita et al. 2011).For example, arabinose-utilization genes are turned on by the AraC regulator in presence of arabinose and cAMP-CRP complex(Johnson and Schleif 1995).Likewiserhamnoseand xylose-utilization genes are induced by RhaR-rhamnose, and XylR-xylose complexes respectively(Egan and Schleif 1993, Song and Park 1997). In addition, some utilization pathways can also be induced by non-specific inducers. For example, rhamnoseutilization genesare known to be induced by structurally similar sugars, L-lyxose or L-mannose(Via, Badia et al. 1996).Similarly, cross induction of genes in fucose metabolic pathway by L-fuculose-1-phosphate, which is a metabolic by product of L-rhamnose hasbeen reported(Chen, Tobin et al. 1987).Carbohydrate transporters in bacteriaare also known to mediate transport of non-cognate substrates(Heller, Lin et al. 1980, Iida, Harayama et al. 1984, Hernandez-Montalvo, Valle et al. 2001).Uptake of xylose by arabinose-transporter AraE provides one such example of cross transport(Hasona, Kim et al. 2004).In addition to cross induction and cross transport, a regulator of one system can also act as a regulator for another(Desai and Rao 2010). For example AraC and XylR, the activators of arabinose and xylose systems respectively inhibit the expression of genes responsible for the other sugar’s utilization (Koirala, Wang et al. 2015). Lastly, bacteria exhibit preferential sugar utilization. In this hierarchy, glucose is, usually, the most preferredsugar, followed by others(Inada, Kimata et al. 1996). This hierarchical utilization of sugars is known as Carbon Catabolite Repression (CCR)(Bruckner and Titgemeyer 2002, Gorke and Stulke 2008)and occursdue to lower levels of the key signalling molecule cAMP which in complex with CRP acts as a positive regulator for induction of most non-PTS sugar regulons(Makman and Sutherland 1965)(b) inhibition of secondary sugar transporters (inducer exclusion) by PTS components(Okada, Ueyama et al. 1981)and (c) post transcriptional regulation by small RNAs(Beisel and Storz 2011).

An additional hierarchy of utilization exists among non-glucose sugars as well. In a recent work, Aidelberg and co-workers studiedsugar utilization in *E. coli*(Aidelberg, Towbin et al. 2014), and demonstrated that these sugars were shown to be consumed in the order: lactose – arabinose - xylose – sorbitol – rhamnose – ribose. Two mechanisms responsible for this hierarchical utilization wereproposed: (a) cross inhibition of utilization genes of sugar systems lower on the preference list by regulators of sugar systems having higher preference and differential activation of sugar systems by cAMP-CRP complex.

The main focus of this study is to explore how short-term adaptation of *E.coli* indefined carbon environments affectsthedynamics of growth and sugar utilization. Specifically we ask (a) does short-term adaptationrewires regulatory circuits in bacteria, specifically withrespect to cross regulatory interactions between specific sugar regulons? And(b) if so, what is the dynamics of this rewiring? To answer these questions, we perform evolution in five different selection conditions, each characterized by presence of a different carbon source. Specifically, the experiments were carried out in environments containingglucose, arabinose, xylose, rhamnose, and a mixture containing all these sugars in a predefined ratio.Among these sugars, glucoseis directly metabolized through glycolysis(Kresge, Simoni et al. 2005). Rhamnose is degraded to rhamnulose-1-phosphate which is cleaved to form dihydroxy acetone phosphate and L-lactaldehyde. The former enters glycolysis and the latter depending on aerobic or anaerobic conversion, gets converted into pyruvate via lactate, a glycolytic intermediate or to propane1,2-diol(Baldoma and Aguilar 1988).On the other hand, pentoses like arabinose and xylose are degradedinto pentose phosphate pathway intermediate xylulose-5-phosphate which is eventually converted into glycolytic intermediates fructose-6-phosphate and glyceraldehyde-3-phosphate(David and Weismeyer 1970, Lawlis, Dennis et al. 1984, Schleif 2000).Although, someof the crosstalk interactions among these systems have been known to exist(Gorke and Stulke 2008),new interactions and their implications have been highlighted in two recent reports(Aidelberg, Towbin et al. 2014, Koirala, Wang et al. 2015).In this context, we are particularly interested to see emergence of changes in dynamics of gene expression, growth, and other physiological changes in a short time frame (∼2000 generations). We presume that changes to the utilization systems will be more “easily” evolved via mutations in regulatory sequences compared to enhancing enzyme activity. Since transcriptional crosstalk in sugar systems mostly involves binding of non-cognate regulator to the upstream regulatory regions of other systems(Desai and Rao 2010), we sequence the upstream regulatory regions of genes in sugar-utilization regulons in the evolved strains to identify mutations responsible for better adaptation.Our results reveal no mutations in these regions and hence indicate that cross talk among sugar-utilization systems are robust and do not, in the framework of our experiment, undergo rewiring in response to sugar selection pressure. However, evolved strains, irrespective of the sugar environment in which they were adapted, show higher fitness than the ancestor in all sugar environments studied in this work(Kassen 2002). All evolved strains show reduced duration of lag phase and increased exponential growth rate compared to the ancestor.However, this enhanced performance of the evolved strains in the carbon environmentsis linked to reduced survival in stationary phase.We were not able to pin point the exact targets where mutations might have taken place, however based on previous reports and the nature of phenotypic changes that were exhibited by the strains, we have commented on targets which might have evolved mutations resulting in the phenotypic changes observed in our study.

## 2. Methodology

### 2.1 Adaptive Laboratory Evolution

We used *E. coli* K-12 MG1655 strain to study the process of adaptive evolution under sugar selection pressures. We propagated this strain under five independent sugar environments - glucose, arabinose, xylose, rhamnose (each individual sugar being present at 0.2% concentration) and lastly, a mixture of all these four sugars.The mixture contained glucose, arabinose, xylose and rhamnose in 6.94: 3.8: 0.66: 0.44 molar ratios, respectively. The final concentration of each of the constituent sugars in the mixture therefore became: glucose – 0.125%, xylose – 0.057%, arabinose – 0.01% and rhamnose-0.008%. Cells were inoculated in 2ml tryptone broth in a test tube containing 0.2% sugars at 37°C with shaking at 160 rpm. Presence of tryptone in the growth medium, may influence the growth of the strains to some extent. However, use of tryptone broth will lead to higher carrying capacity than minimal media. This may allow greater supply of beneficial mutationsas the population bottle-neck will contain more cells during transfer into a fresh medium.

Evolution in a given sugar medium was carried out by sub-culturing 1:100 of the overnight culture into fresh medium twice daily. The experiment wascontinued for 2,000 generations. For each sugar environment, two parallel populations were evolved (designated strain 1 andstrain 2), yielding a total of ten different strains for five different sugar environments. The strains were named according to the sugar environment in which they were evolved - Glu1 and Glu2 for the duplicate strains adapted in glucose. Likewise Ara1 and Ara2, Xyl 1and Xyl2, Rha1 and Rha2 and Mix1 and Mix2 for strains evolved under arabinose, xylose, rhamnose and mixed-sugar conditions respectively. Approximately after every 50 generations, strains were stored at -80 degrees for future analysis. The term “evolved strains” has been used to indicate strains evolved for 2,000 generations, unless otherwise mentioned.

### 2.2 Phenotypic Characterization

To understand the phenotypic changes occurring in the strains under diverse sugar selection pressures, we measured three parameters associated with the bacterial growth in the evolved strains, and compared them with those in the wild-type *E. coli.* The three parameters measured were as follows: (a) duration of lag phase upon sub-culture, (b) growth rate in the log-phase, and (c) steady state optical density (OD) values. These parameters were measured in strains evolved for300, 600, 1,200 and 2,000 generations. For each strain, these measurements were made in all five sugar environments.To make these measurements, single colonies of ancestor and evolved strains were inoculated into tryptone broth and grown till saturation. Next, they were sub-cultured 1:100 and grown in the same medium containing the appropriate sugar in a micro plate reader with shaking at 37°C and optical density (OD600) readings taken every 45 minutes for 16 hours. The duration of lag phase was calculated as previously described(Zwietering, Rombouts et al. 1992). Growth rate of the strains were calculated as (*ln*0D2 - *ln*0D1)/(t2 – t1), where OD1 and OD2 were taken to be theOD600 values at time t1 and t2 respectively in the exponential growth curve. The times t1 and t2 weretaken to be the time points when the exponential growth phase begins and ends respectively. Steady state OD600 values for different strains were taken to be the respective OD600 value at which growth becamesaturated. Each experiment was performed in triplicate and average values with standard deviation reported.

### 2.3 DNA Sequencing

The promoter regions of*ptsG*(for glucose utilization system), *araE*, *araFGH*, and *araC*-*araBAD*(for arabinose utilization system), *xylE* and *xylFGH*(for xylose utilization system) and *rhaBAD* and *rhaT*(for rhamnose utilization system) were sequenced in ancestor and evolved strains. All sequencing reactions were performed by Eurofins Bangalore by using cycle sequencing technology on ABI 3730XL sequencing machines.Prior to sending out samples for sequencing, DNA from single colonies was PCR amplified using primers as specified in Table S1. The purity and concentrations of DNA samples weredetermined using bio spectrometer from Eppendorf. Samples showing260/280 absorbance ratio between 1.6 – 2were considered pure and taken for sequencing.Each DNA strand was sequenced at least twice before confirming mutation (if any). Multiple sequence alignment for each regulatory region of the sugar utilization genes in ancestor and evolved strains was done using a T-coffee alignment tool(Notredame, Higgins et al. 2000).

### 2.4 Cell Survival in Stationary Phase

Ancestor and evolved strains were inoculated from overnight cultures into fresh tryptone broth containing the sugar in which the strains were adapted and thereafter, incubated at 37°C and grown for about 24 hours. After incubation, samples were collected and appropriate dilutions plated on LB agar plates. The number of colonies in each plate was counted and was reported as Colony Forming Units per ml (CFU/ml). This indicated the proportion of population surviving in stationary phase.

### 2.5 Reporter Plasmids

Gene expression measurements for *araBAD, xylAB, rhaBAD,* and *ptsG* were prepared using the CRIM plasmid pLA2, where the *lacZ* was replaced by *gfp*(Haldimann and Wanner 2001).Five hundred bases upstream of the first gene in the respective operons were cloned in the plasmid and promoter activity quantified as fluorescence (A.U.). For measuring promoter activity, evolved strains were grown in tryptone broth containing 0.2% of the respective sugar (in which they were adapted), and expression was measured when the cells reached mid-log phase of growth. Prior to expression measurements, evolved strains were pre-cultured in adaptation environment for at least two successive transfers. The gene expression level in each evolved strain was compared to that of the ancestor, grown on the same sugar. The expression of reporter plasmids in absence of any sugar was taken as negative control.

### 2.6 Statistical analysis

Student’s t-test with equal variances was used to quantify significance of observed differences between ancestral and evolved strains, where* indicates p<0.05, ** indicates p<0.005, and ns indicates not significant.

## 3. Results

### 3.1 E. coli strains evolve towards shorter lag phases and higher growth rates in the selection environment

Using adaptive laboratory evolution we tried to understand the physiological and genetic adaptation of *E. coli* grown in defined sugar environments. In these experiments, cells were adapted in five different sugar environments, namely glucose, arabinose, xylose, rhamnose, and the fifth one consistingof a mixture of all four sugars. Starting from an ancestor *E. coli* strain (the ancestral strain), two parallel populations were evolved in each sugar environment. After 2,000 generations in a given selection environment, evolved strains show reduced lag duration and increased growth rate compared to the ancestral strain.

During the course of the experiment, as cells were repeatedly sub-cultured into the same medium, they presumably evolved towards maintaining ahigherbasal level of transporters and enzymes required for utilization of a particular sugar. Unlike the ancestor, the evolved strains could therefore initiateutilizationof the sugar faster upon transfer into fresh medium. This eventually also led to faster carbon source utilization; consequently higher growth rates in evolved strains compared to the ancestor. In most cases, thefinal cell concentrations remained same in ancestral and evolved strains leading to comparable steady state OD600 values. However, few evolved strains showed higher steady state OD600 values than the ancestor indicating higher biomass yield (Figure 1B, 2A-B, S1B, S2B, and S4B). This suggested that while the dynamics of sugar utilization in the bacterium were tunable within a short evolutionary timeframe, the efficiency of utilization of a carbon source did not change in a relatively brief evolutionary experiment in most cases.

**Figure 1.**
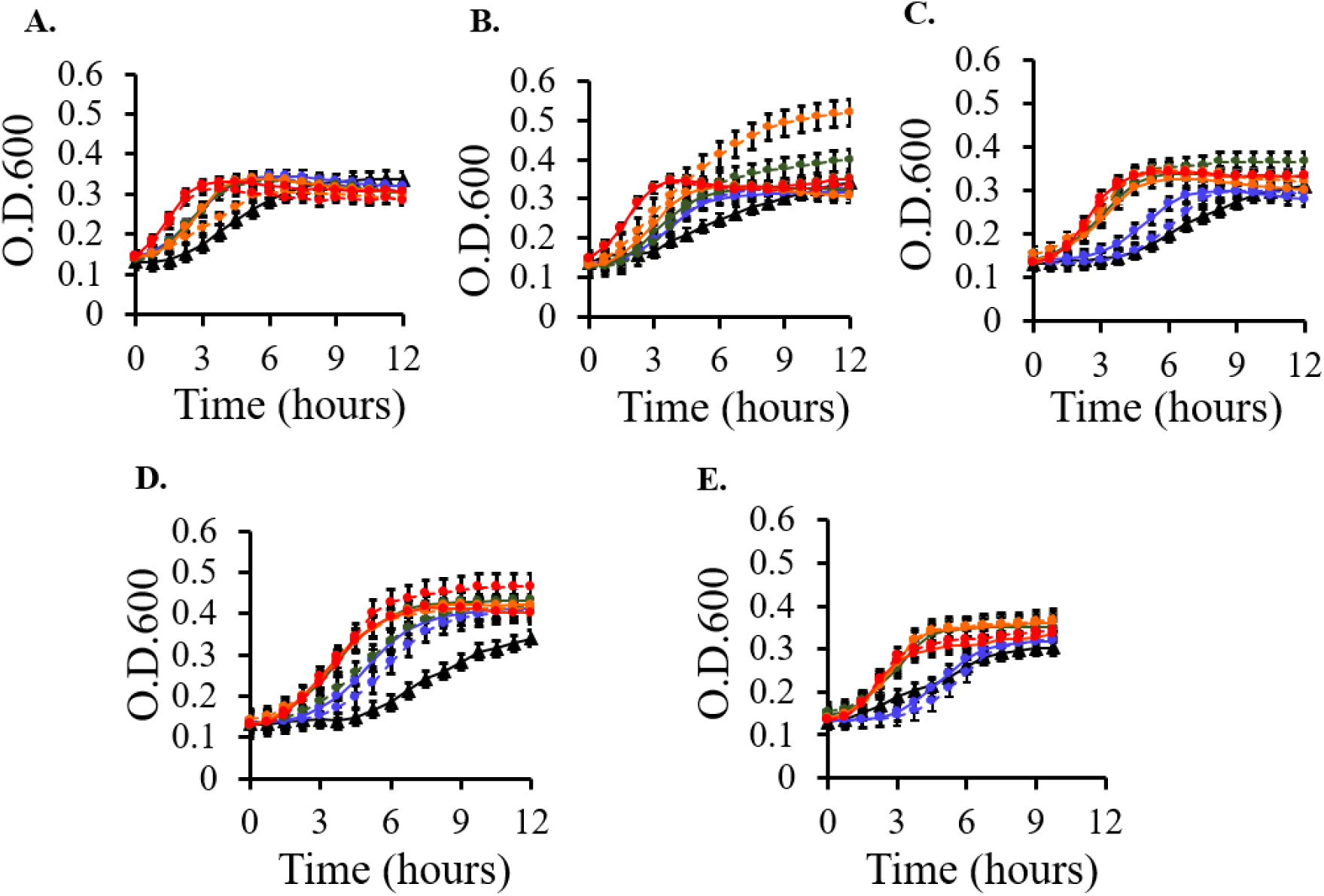
Dynamics of growth of ancestral and evolved strains 1 & 2 from different time points in selection environments: A. Glu1 & Glu2 in glucoseB. Ara1 & Ara2 in arabinose C. Xyl1 & Xyl2 in xylose and D. Rha1 & Rha2 in rhamnose and E. Mix1 & Mix2 in sugar-mixtures. Evolved strain 1 has been shown by solid lines and strain 2 by dashed lines. The different time points are indicated by colour coding:blue (200^th^ generation), green (600^th^ generation), orange (1200^th^ generation) and red (2000^th^ generation). Ancestral strain is indicated by black solid line. Data indicate average values for three independent experiments. Error bars denote standard deviation from three separate experiments performed on different days.

Next, we wanted to see the time of appearance of the acquired traits in evolved strains in different selection environments. To do this, we measured lag phase length and growth rate in both the evolved strains from 300, 600, 1200, and 2,000 generations in all environments and compared with the ancestral strain. We also performed statistical analysis to test the level of significance of the observed differences between evolved and ancestral strains. Table 1. summarizes the mean lag phase length, growth rate, and steady state OD600 data for all strains from different time points in different selection environments. The p-values indicated for each parameter (lag phase or growth rate) in evolved strains have been calculated with respect to the ancestor. Figure1 A-E shows the corresponding growth curves for evolved and ancestral strains.In glucose and arabinose evolved strains, both the acquired traits appeared by 300 generations. For glucose evolved strains, the reduction in lag phase was achieved in a single step, while in arabinose evolved strain, lag phase reduced in two steps. Growth, increased gradually over 2,000 generations for arabinose evolved strains, while for glucose evolved strains, increase in growth rate was not significant after 1,200 and 600 generations for strain 1 and 2 respectively. For both xylose and rhamnose evolved strains, fold reduction in lag phase increased by 600 generations. Growth rate increased slowly and irregularly over 2,000 generations in both the evolved strains.In strains evolved in sugar-mixtures, lag phase in ancestor and evolved strains did not vary significantly till 1,200 generations. However, in both the evolved strains, growth rate increased progressively with time.

**Table 1.**
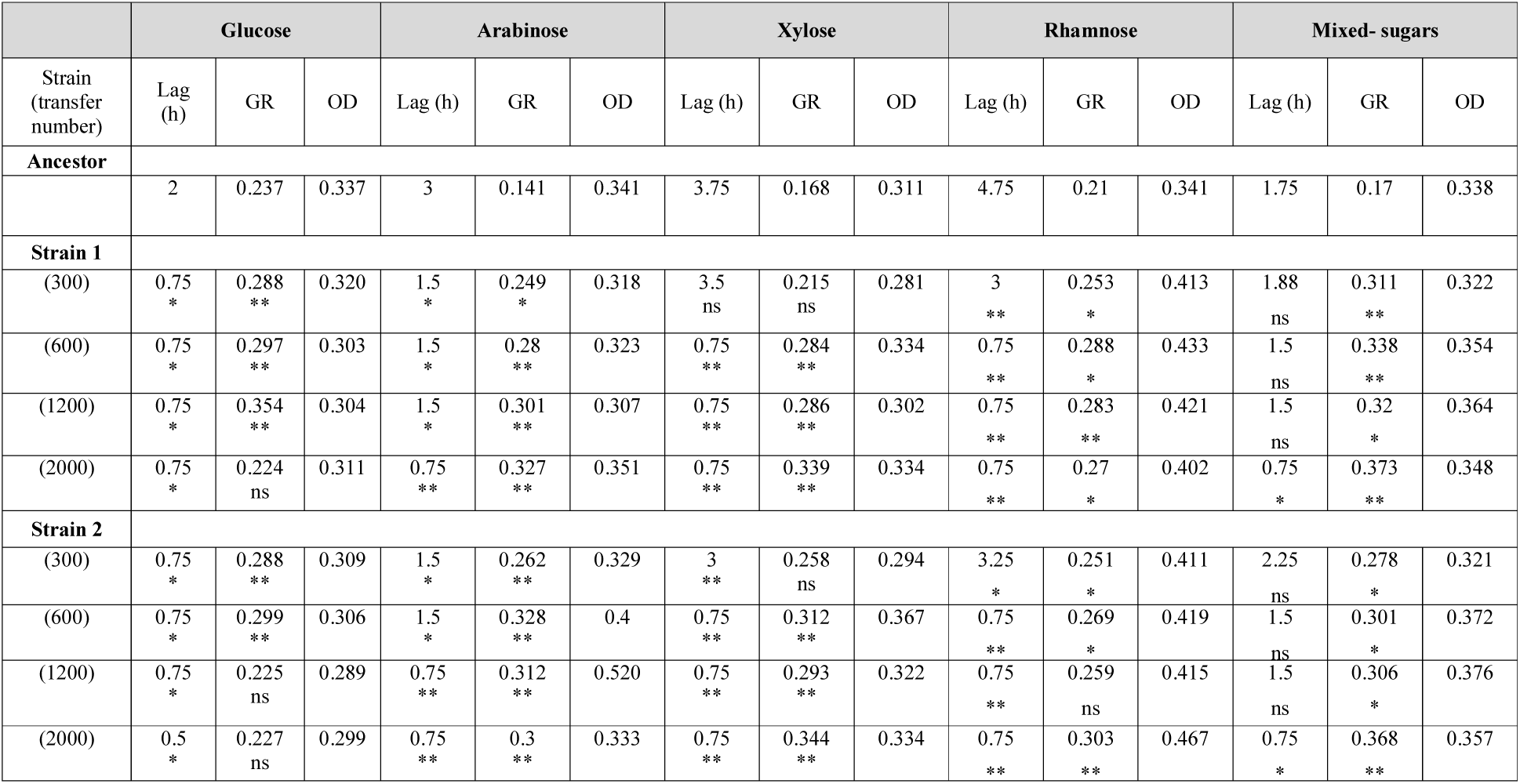
Lag phase duration, growth rate and steady state O.D. 600 values for ancestor and evolved strains from different time points in selection environments

### 3.2 Increase in growth rate and reduction in the duration of lag-phase is independent of environment

Next, we wanted to study the growth characteristics of the evolved strains in other sugar environments in which they were not evolved.Similar to adaptation environments, the evolved strains also exhibited reduced lag, increased growth rate and comparable steady state OD600 values with respect to the ancestral strain (Figures 2 A-D, S1-4, and Table S2A-E).

**Figure 2.**
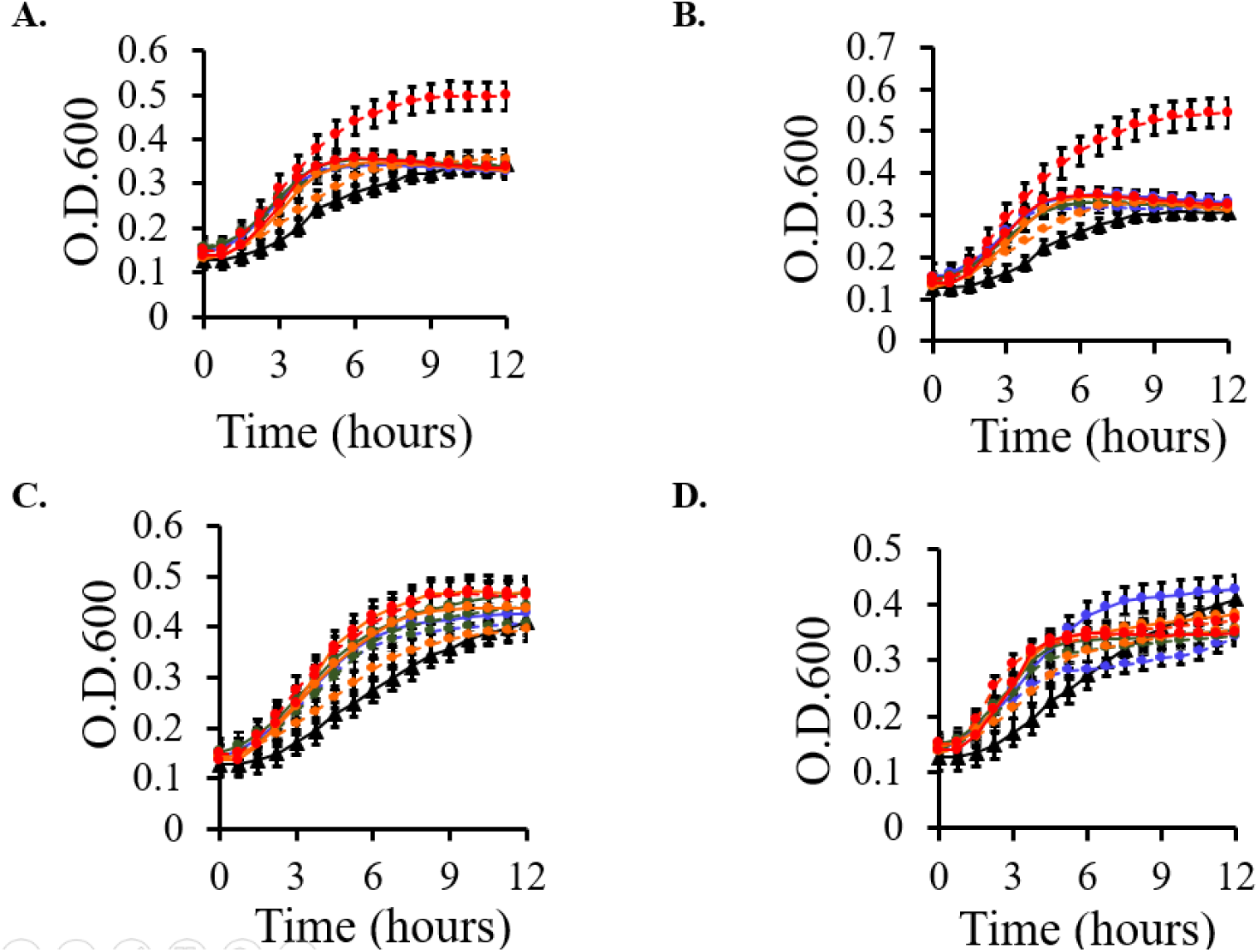
Dynamics of growth of ancestral and glucose evolved strains from different time points in non-selection environments: A. Arabinose B. Xylose C. Rhamnose and D. Mixed-sugars. Evolved strain 1 has been shown by solid lines and strain 2 by dashed lines. The different time points are indicated by colour coding: blue (200^th^ generation), green (600^th^ generation), orange (1200^th^ generation) and red (2000^th^ generation). The ancestral strain has been indicated by black solid line. Data indicate average values for three independent experiments. Error bars denote standard deviation from three separate experiments performed on different days.

Since theevolved strains, irrespective of the environment of adaptation, showed improved growth characteristics across non-selection environments, we explored dynamics of fixation of the acquired traits in the evolved strains in non-selection environments as well. Table.S2A-E summarizes the data for lag phase length, and growth rate in evolved strains in different non-selection environment and indicates the significance of difference of these parameters from the ancestral strain. In glucose evolved strains, for example, the dynamics of fixation of lag phase trait across different sugar environments was similar to that observed in selection environment. However, increase in growth rate trait appeared late (in arabinose) or changed inconsistently over the given period of evolution (in xylose) or was not significantly different from that of ancestor (in rhamnose). In glucose evolved strains grown in sugar-mixtures, growth rate increased irregularlyfrom 300 generations and continued till 2,000 generations.In arabinose evolved strains, both the acquired traits appeared by 300 generations in all environments except glucose where the reduced lag phase trait was fixed much later. In xylose and rhamnose evolved strains, the dynamics of fixation of the two traits was similar across selection as well as non-selection environments. For mixed evolved strains, the fixation of lag phase occurred later in other sugar environments, compared to its selection environment.

### 3.3 Lag phase optimization and increased growth rate were not observed because of mutations in the upstream regulatory elements of the sugar utilization genes

The evolved strains performed better than the ancestor by (a) reducing the duration of lag phase and (b) increasing their growth rate. Can acquired phenotypes of the evolved strains be attributed to increased expression levels of the transport and/or utilization genes? Since dynamics of expression of the sugar utilization genes were dictated largely by the architecture of the regulatory region, we sequenced the upstream regulatory regions of sugar transporters and metabolic genes. We were particularly interested to see if there is any change in DNA sequences in the sites where regulatory proteins or RNA polymerase binds, -35 or -10 elements, or the transcription start site. We looked for mutations in eight upstream regulatory regions of the following genes - *ptsG*, *araE*, *araFGH*, *araC*-*araBAD*, *xylE*, *xylFGH, rhaBAD* and *rhaT*.

None of the upstream regulatory regions show any mutation except for *rhaT* (Figure 3 and S5-S10). Promoter region of *rhaT*gene showed mutation in the CRP binding site in three of the ten evolved strains. One of the glucose-, rhamnose-, and mixed-sugar-evolved strains showed insertion of a single nucleotide in the CRP binding site. For Mix1 strain, the insertion was at position -95, while that in Glucose1 and Rhamnose1 strain the location of insertion was at -90 from the transcriptional start site. This indicates strong selection in this locus. The next question that follows from here is whether this mutation offers any advantage to bacteria? Specifically, is this mutation responsible for reduced lag and increased growth rate in Glucose1, Rhamnose1, and Mix1 strains grown in rhamnose and other sugars?Length of lag phase and growth rate in these strainswas observed to be comparable to other evolved strainsindicating that this mutation may not contribute directly to establishment of the two traits. Investigating the physiological impact of this mutation on evolved strains will require further experimentation. Absence of mutation in other regulatory regions may indicate that both a) these loci are not strong targets of positive selection or b) appearance and establishment of mutation in these loci is slower than that of *rhaT*. In fact, in a long term evolutionary experiment, dynamics of appearance of mutations in different targets vary (Schneider et.al, *Genetics*, 2000).

**Figure 3.**
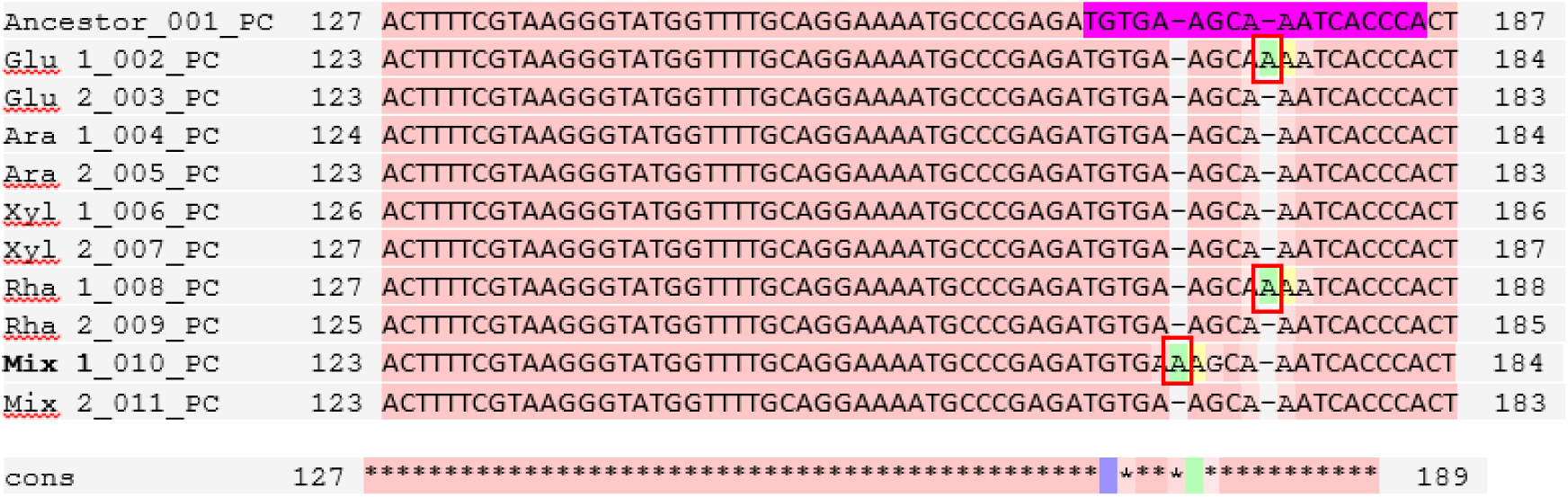
Sequence alignment data of ancestor and evolved strains showing the CRP binding site in *rhaT* promoter (highlighted in pink). Glucose 1, Rha1 and Mix 1 strains show single nucleotide insertion in this region (inserted bases highlighted by red boxes). *rhaT* regulatory region was sequenced using cycle sequencing technology by Eurofins Bangalore and multiple sequence alignment was done using T-coffee online tool.

### 3.4 Reduced lag phase in evolved strains is associated with reduced survival in the stationary phase

The evolved strainsfrom our experiments exhibited a considerablyshorter lag period compared to the ancestor. This is likely due to presence of transporter and utilization proteins maintained in cells during stationary phase, which could initiate immediate utilization of the sugar upon transfer to fresh medium(Madar, Dekel et al. 2013, Gefen, Fridman et al. 2014).However, maintenance of these extra proteins in the stationary phase of the cells must incur some cost to the cell.This cost may be associated with reduced survival of the evolved strains in the stationary phasecompared to the ancestor. To test this possibility, we compared the CFUs in the wild-type and the evolved strain when grown for an extended period in stationary phase. In adaptation environments, evolved strains were shown to form fewer colonies than the ancestor and exhibited approximately 2-5 fold reduction in colony forming unit/ml than the ancestor (Table 2 and Figure S11). Therefore, the cost of maintaining extra proteins in the stationary phase by the evolved strains was accompanied by increased cell death. The ancestral strain, on the other hand, did not have these stored proteins and needed to synthesize them prior to beginning active growth; leading to longer lag.Similar results wereobtainedby Oxman and co-workerswith *E. coli* strains adapted in succinate minimal media for a few hundred generations(Oxman, Alon et al. 2008).

**Table 2.** Comparison of CFU/ml of ancestral and evolved strains in adaptation environment

| Strain | Environment | CFU/ml | Fold reduction compared to the ancestor |
| --- | --- | --- | --- |
| Ancestor | Arabinose | 35000 +/- 2200 | 5 |
| Evolved |  | 7000 +/- 800 |  |
| Ancestor | Glucose | 5200 +/- 248 | 1.9 |
| Evolved |  | 2800 +/- 141 |  |
| Ancestor | Xylose | 9000 +/- 4250 | 1.8 |
| Evolved |  | 5000 +/- 1178 |  |
| Ancestor | Rhamnose | 36000 +/- 1400 | 2.4 |
| Evolved |  | 15000 +/- 2500 |  |

### 3.5 Short-term evolutionary experiment does not alter regulatory crosstalk between sugar utilizations systems

One of the questions of interest in this study was to discover to what extent does short term evolution alters cross-regulatory interactions between sugar utilization regulons. Hence, we were particularly interested in studying, for instance, how arabinose mediated repression of xylose change when cells are evolved in xylose? To answer these questions, we used a single-copy plasmid with *gfp* as a proxy for gene expression to analyse the extent of regulatory crosstalk between sugar utilization systems.

Our results show that while evolution in defined carbon environments does not qualitatively alter the regulatory pattern associated with the different sugar utilization systems; there are distinct quantitative changes in the expression patterns. As shown in Figure 4, two distinct features of this phenomenon are apparent. First, during the course of the evolutionary experiments, the activity of the promoter associated with the sugar in which the bacteria were grown has increased by upto 50%. This is consistent for all sugars, and both repeats for each environment. Second, we also note a marginal decrease (not statistically significant, data not shown) in the repression associated with a sugar system over another sugar system. For instance, when evolved in xylose, the extent to which the xylose system is repressed by arabinose is slightly lower in the evolved strain when compared to the wild type.

**Figure 4.**
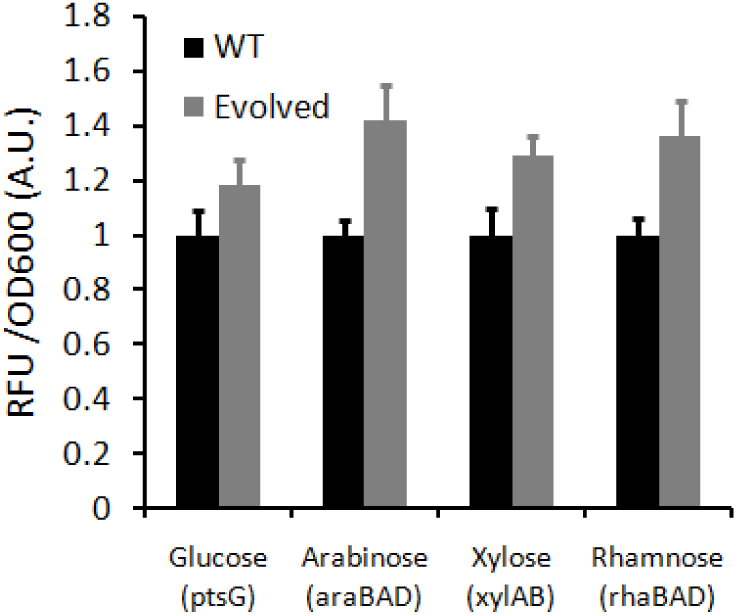
Expression of sugar utilization genes in wild-type and evolved strains. All evolved strains exhibit an elevated expression level of the sugar locus in which they were evolved. Wild-type expression levels in each sugar are normalized to one.

## 4. Discussion

In this work, we aim to understand the nature and dynamics of short-term adaptation of bacterial strains to sugar selection pressures. Starting from an ancestral *E. coli* strain, evolutionary experiment was carried out in each of the five different sugar environments namely glucose, arabinose, xylose, and rhamnose, and in sugar-mixtures for 2,000 generations. All evolved strains exhibited a lower lag phase and higher a growth rate as compared to the ancestor. This change in growth dynamics in the evolved strainswas independent of the sugar environment in which the experiment was conducted, and was therefore, independent of genetic sequence at the locus associated with utilization of a precise sugar. We also noted that a short-term adaptation does not (a) change the qualitative pattern of regulatory interactions between these sugar utilization systems, and that (b) no changes in the regulatory sequence take place for any of the genes involved. Moreover, the hierarchy associated with *E. coli* using these sugars is strictly maintained in all evolved strains. Nevertheless, we see quantitative changes in the expression levels of different sugar utilization genes in the evolved strains. More precisely, the expression levels for the genes involved in utilization of the carbon source in which the bacteria were evolved were increased; and all other sugars which the bacteria did not “see” for 2000 generations were slightly repressed. In addition to reduced lag phase and increased growth rate phenotypes, few evolved strains also exhibited increased biomass yield. Although, trade-off between growth rate and biomass yield have been reported earlier, we did not observe such trade-off in this study (Novak, Pfeiffer et al. 2006).

Based on previous reports, we also tried to predict the possible targets which might have been subjected to positive selection during the course of our evolution experiment. Mutations in a number of candidate genes were reported to be responsible for reduced lag phase, increased growth rate, larger cell size, improved sugar uptake in strains subjected to adaptation under glucose limiting conditions(Schneider, Duperchy et al. 2000, Maddamsetti, Hatcher et al. 2017). Of these, mutation in *spoT*, a gene whose product regulates stringent response was shown to result in both reduction in lag phase as well as increase in exponential growth rate. Mutation in another gene*nadR*, coding for a repressor of genes involved in NAD metabolism was presumed to be responsible for increased growth rate or reduced lag phase or both in evolved strains (Schneider, Duperchy et al. 2000). Lastly, mutations in *topA* and *fis* genes was reported to be responsible for increased growth rate in evolved strains. Mutations in several other genes imparted other improved traits in evolved strains – for example strains having mutation in *pykF* showed improved glucose uptake, or in *pbpA*-*rodA* operon resulted in larger cell volumes (Woods, Schneider et al. 2006). The glucose evolved strains also showed improved performance in environments containing those substrates which shared similar mechanism of transport as that of glucose (Lenski, Mongold et al. 1998). Moreover, increase in biomass yield in strains evolved in glucose was reported to be due to mutations in the glucose transporter genes(Bachmann, Fischlechner et al. 2013).

In our experiments, the potential targets for positive selection might be some components common to the utilization pathways of the sugars used. This is most likely because improved growth phenotypes were exhibited by evolved strains across all sugar environments. Among these, glucose is transported via the PTS systems and rest by non-PTS transporters, specific for each pentose(Luo, Zhang et al. 2014). Although promiscuous transporters do exist for pentoses(Khankal, Chin et al. 2008),glucose is not transported through these. Hence sugar transporters seem unlikely as candidates for acquiring beneficial mutations. The next probable targets can be genes present in the metabolic pathways of the hexose and pentose sugars. While glucose enters central metabolism directly through glycolysis, the pentoses are first converted to pentose phosphate pathway intermediate before entering the glycolytic pathway (Mayer and Boos 2005).Hence genes for enzymes catalysing reactions in the latter half of the glycolytic pathway may be targets of positive selection. Potential candidates are*pyfK* and *pyfA*, coding for pyruvate kinases. Another possible target can be a common regulatory gene that controls expression of all these sugar regulons. For example *crp*, the gene coding for the global positive regulator of different sugar regulons can be a target for positive selection. A beneficial mutation in *crp* for example, might lead to increased expression of the target genes which includes sugar transporters, metabolic genes or regulators. This might result in faster utilization of sugar leading to improved growth dynamics. Another target for positive selection can also be *cyaA* gene coding for adenylate cyclase, which regulates the amount of CRP-cAMP in the cell.

From the perspective of cross-regulatory interactions, our results show that in a relatively short time of 2,000 generations, there is likely no significant advantage in the bacterium in losing its ability to utilize carbon sources it is not exposed to. This is likely an important result from the perspective of bacterial evolution in their ecological niches, where the local environments are likely to be fluctuating with time and space(Winfield and Groisman 2003, van Elsas, Semenov et al. 2011).

## Supplementary Material

**Figure S1:**
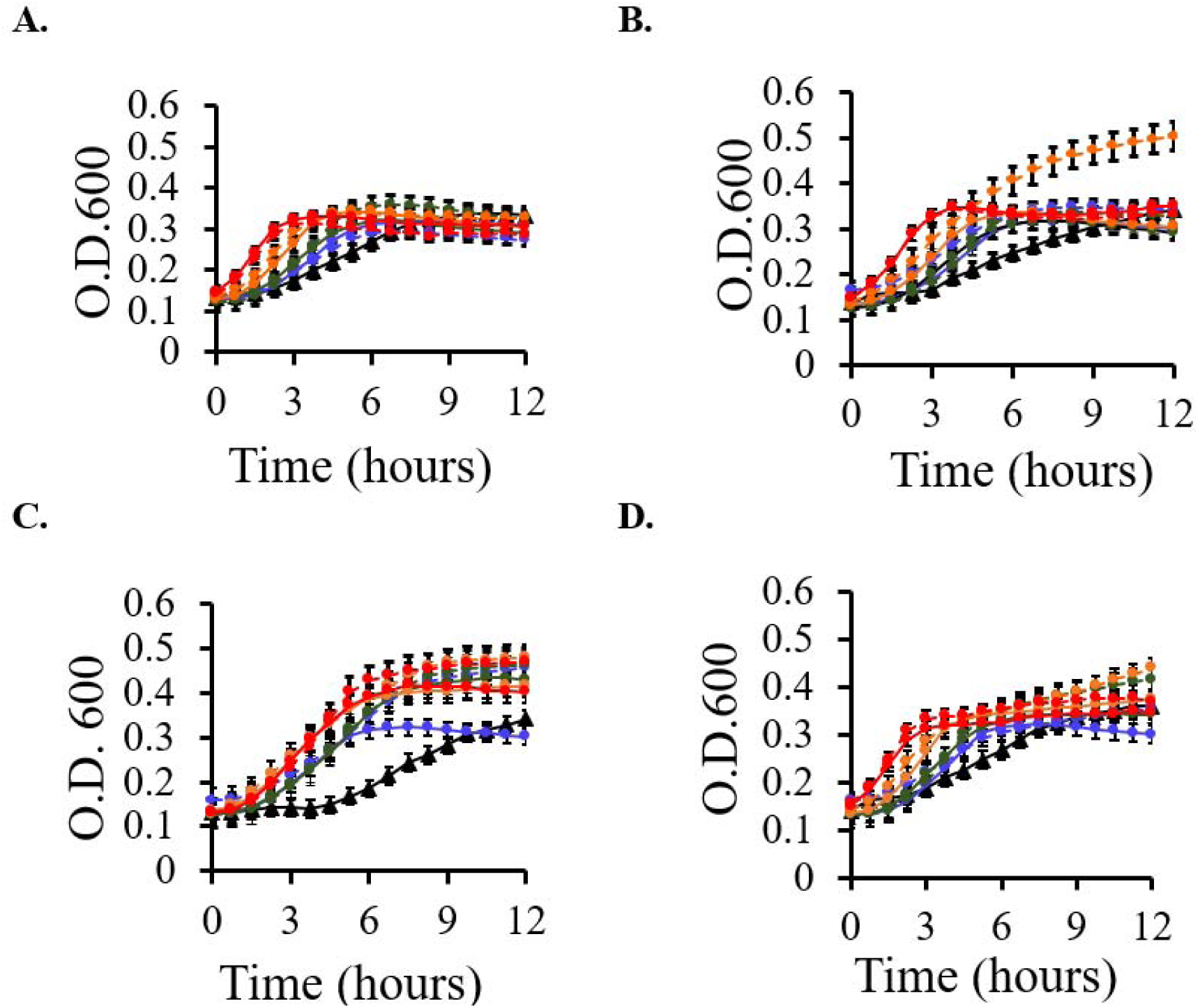
Dynamics of growth of ancestral and arabinose evolved strains from different time points in non-adapted environments: A. Glucose B. Xylose and C. Rhamnose and D. Mixed-sugars. Evolved strain 1 has been shown by solid lines and strain 2 by dashed lines. The different time points are indicated by colour coding: blue (200^th^ generation), green (600^th^ generation), orange (1200^th^ generation) and red (2000^th^ generation). Data indicate average values for three independent experiments. Error bars denote standard deviation from three separate experiments performed on different days.

**Figure S2:**
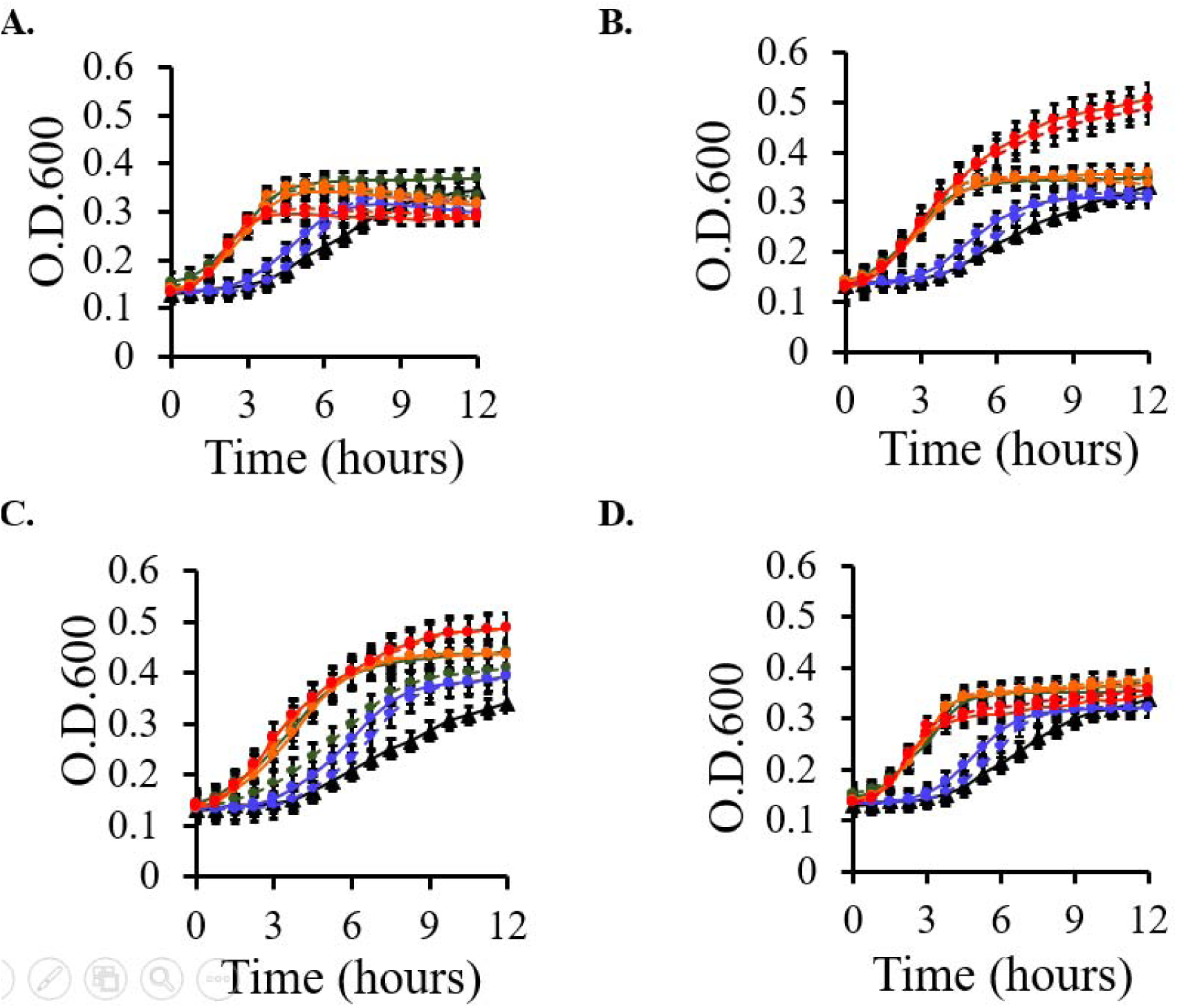
Dynamics of growth of ancestral and xylose evolved strains from different time points in non-adapted environments: A. Glucose B. Arabinose and C. Rhamnose and D. Mixed-sugars. Evolved strain 1 has been shown by solid lines and strain 2 by dashed lines. The different time points are indicated by colour coding: blue (200^th^ generation), green (600^th^ generation), orange (1200^th^ generation) and red (2000^th^ generation). Data indicate average values for three independent experiments. Error bars denote standard deviation from three separate experiments performed on different days.

**Figure S3:**
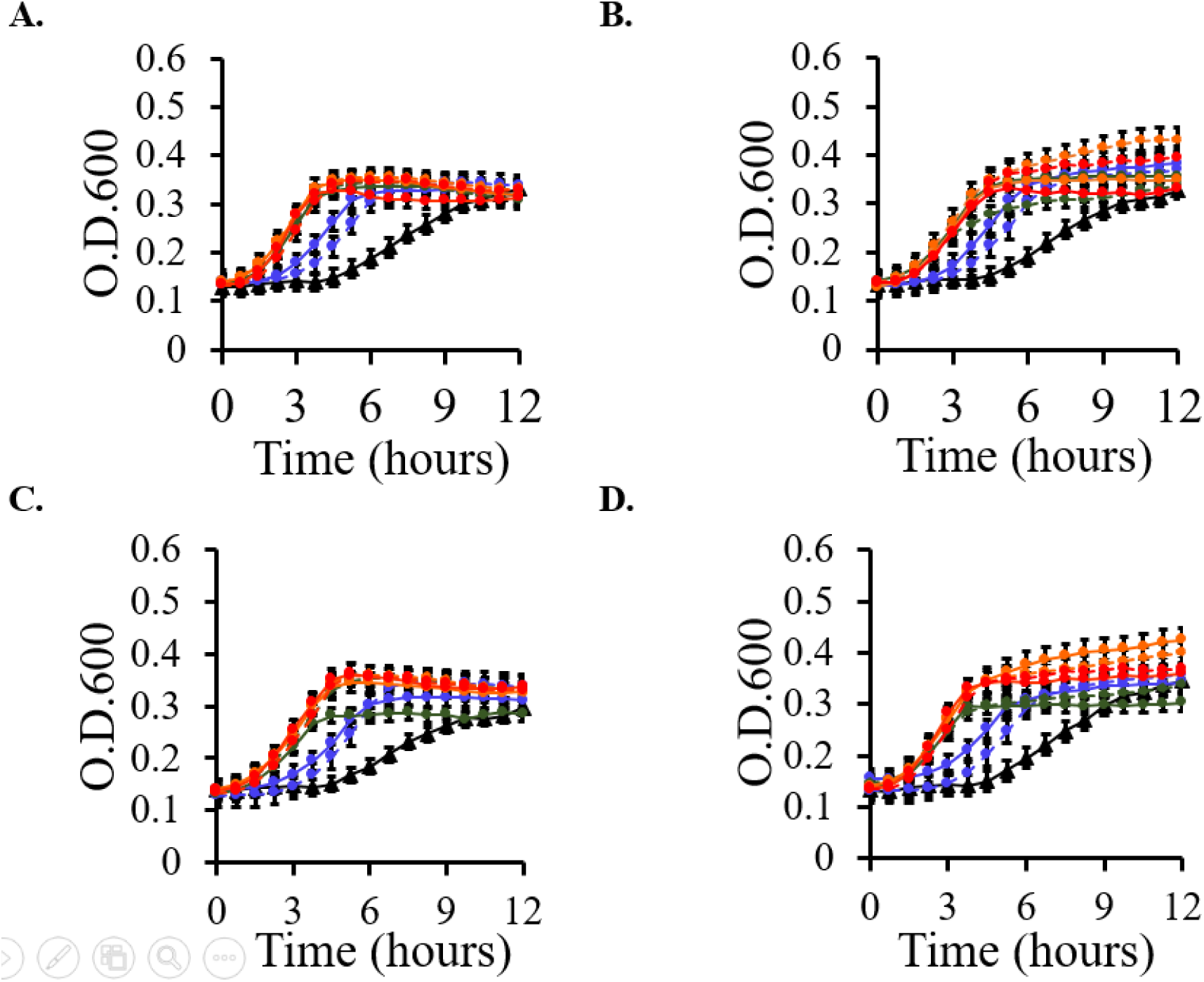
Dynamics of growth of ancestral and rhamnose evolved strains from different time points in non-adapted environments: A. Glucose B. Arabinose and C. Xylose and D. Mixed-sugars. Evolved strain 1 has been shown by solid lines and strain 2 by dashed lines. The different time points are indicated by colour coding: blue (200^th^ generation), green (600^th^ generation), orange (1200^th^ generation) and red (2000^th^ generation). Data indicate average values for three independent experiments. Error bars denote standard deviation from three separate experiments performed on different days.

**Figure S4:**
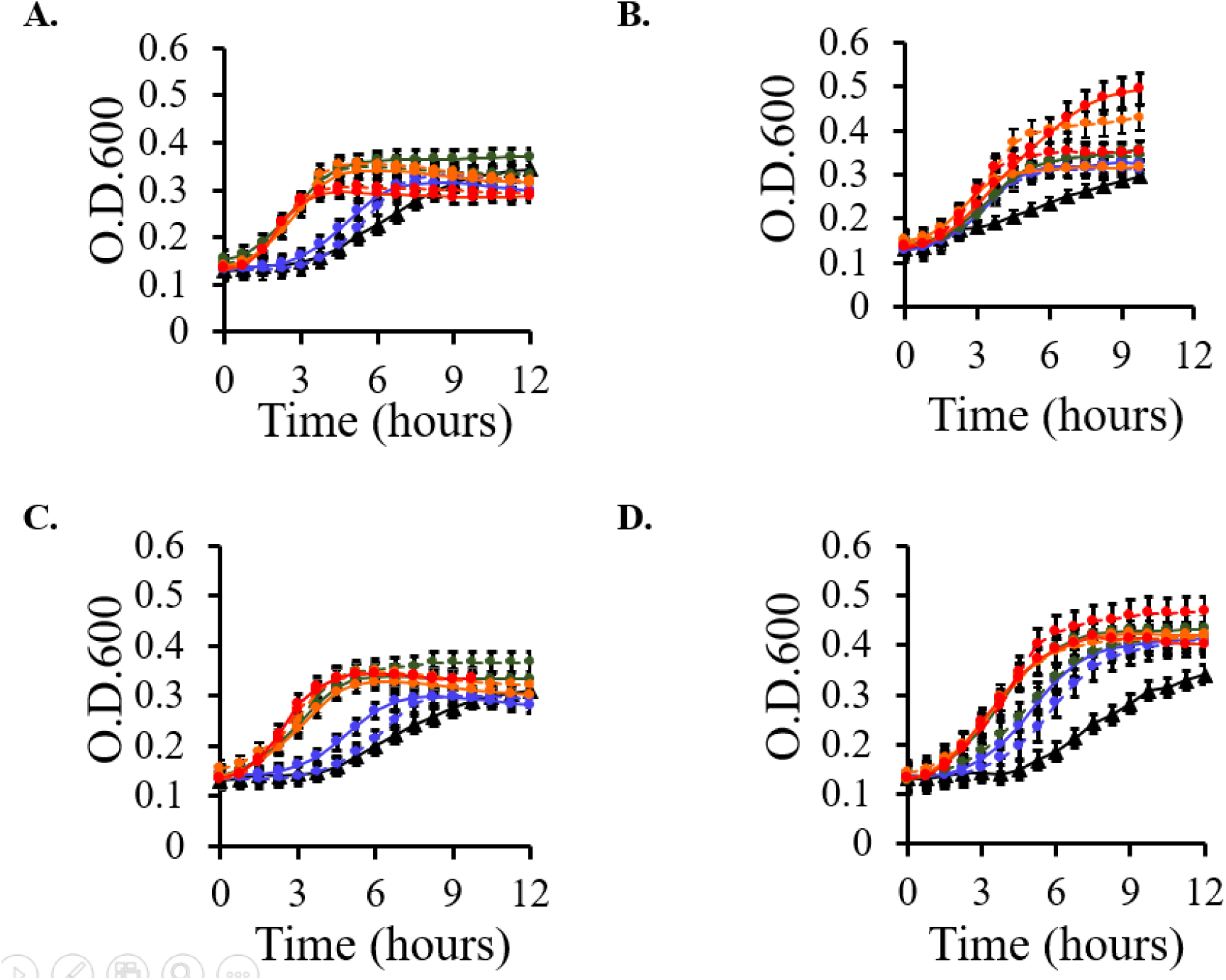
Dynamics of growth of ancestral and sugar-mixture evolved strains from different time points in non-adapted environments: A. Glucose B. Arabinose and C. Xylose and D. Rhamnose. Evolved strain 1 has been shown by solid lines and strain 2 by dashed lines. The different time points are indicated by colour coding: blue (200^th^ generation), green (600^th^ generation), orange (1200^th^ generation) and red (2000^th^ generation). Data indicate average values for three independent experiments. Error bars denote standard deviation from three separate experiments performed on different days.

**Figure S5:**
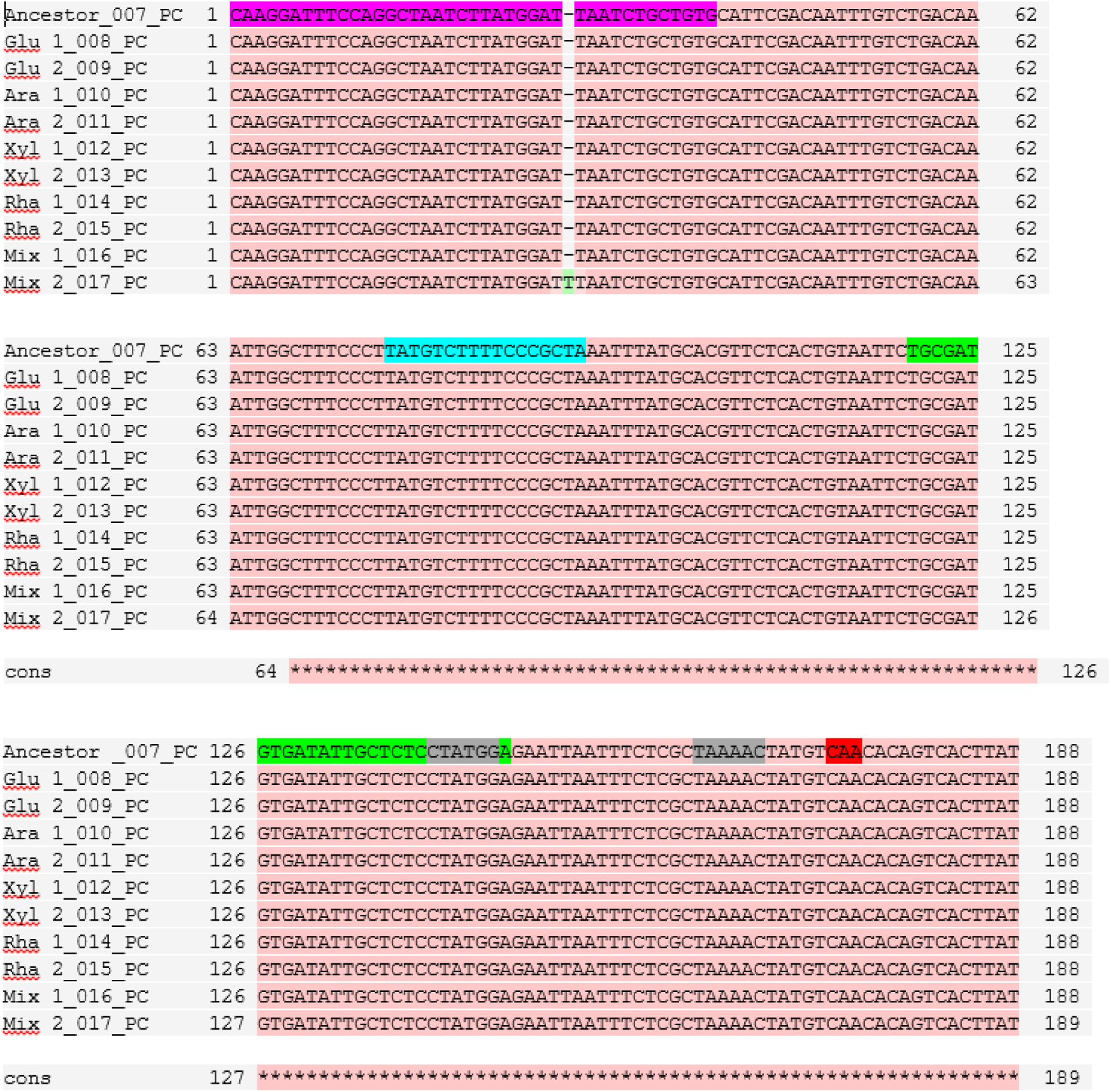
Sequence alignment data of *araFGH* regulatory region in ancestor and evolved strains showing the AraC binding site (pink & blue) CRP binding site (green), -10 & -35 (grey) and transcription start site (red). *araFGH* regulatory region was sequenced using cycle sequencing technology by Eurofins Bangalore and multiple sequence alignment was done using T-coffee online tool.

**Figure S6:**
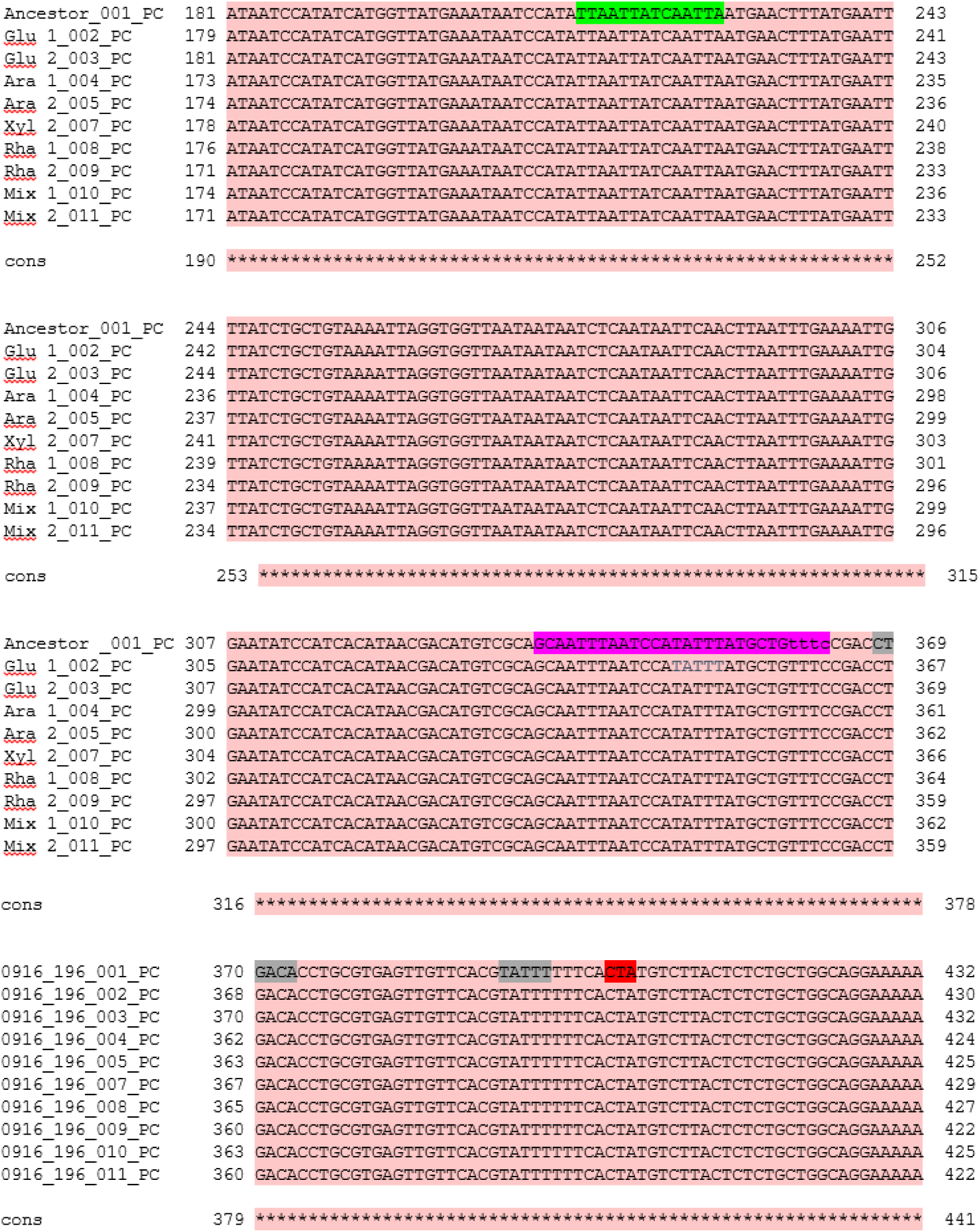
Sequence alignment data of *araE* regulatory region in ancestor and evolved strains showing the AraC binding site (pink & blue) CRP binding site (green), -10 & -35 (grey) and transcription start site (red). *araFGH* regulatory region was sequenced using cycle sequencing technology by Eurofins Bangalore and multiple sequence alignment was done using T-coffee online tool.

**Figure S7:**
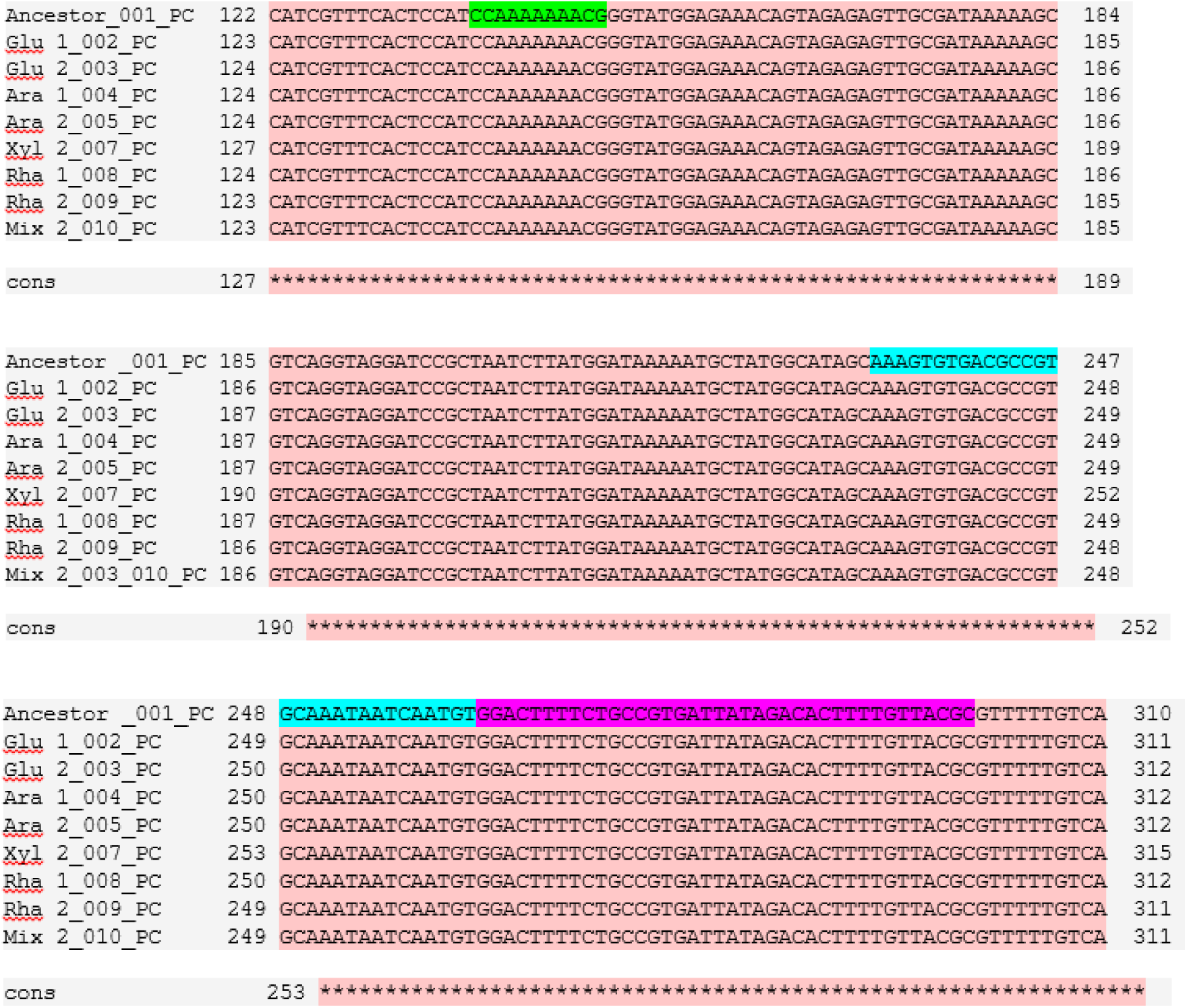
Sequence alignment data of *araC*-*araBAD* regulatory region in ancestor and evolved strains showing the polymerase binding site in *araB* promoter (pink), polymerase binding site in *araC* promoter (green) and CRP binding site (blue). *araC*-*araBAD* regulatory region was sequenced using cycle sequencing technology by Eurofins Bangalore and multiple sequence alignment was done using T-coffee online tool.

**Figure S8:**
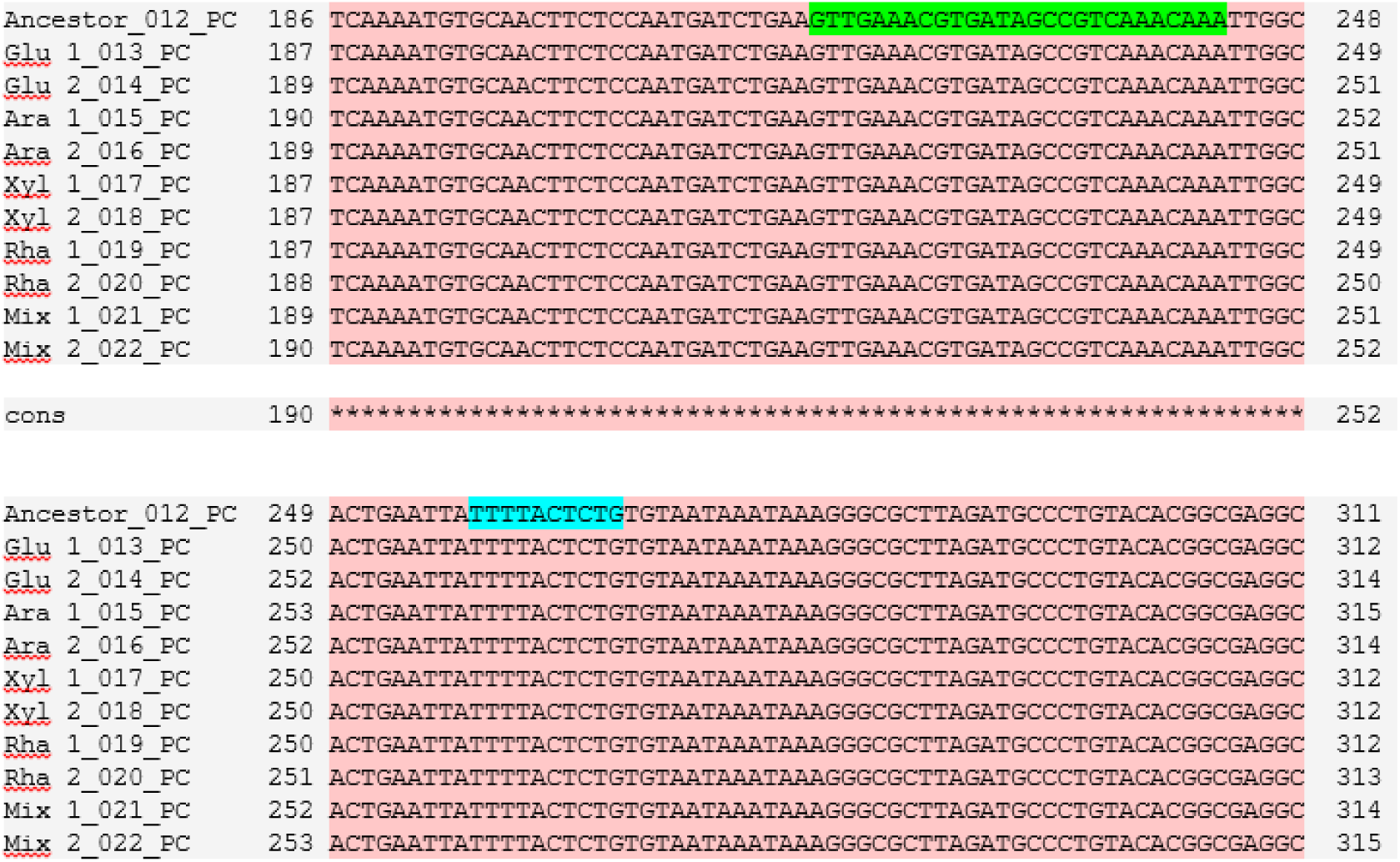
Sequence alignment data of *ptsG* regulatory region in ancestor and evolved strains showing the CRP binding site (green) and Mlc binding site in (blue). *ptsG* regulatory region was sequenced using cycle sequencing technology by Eurofins Bangalore and multiple sequence alignment was done using T-coffee online tool.

**Figure S9:**
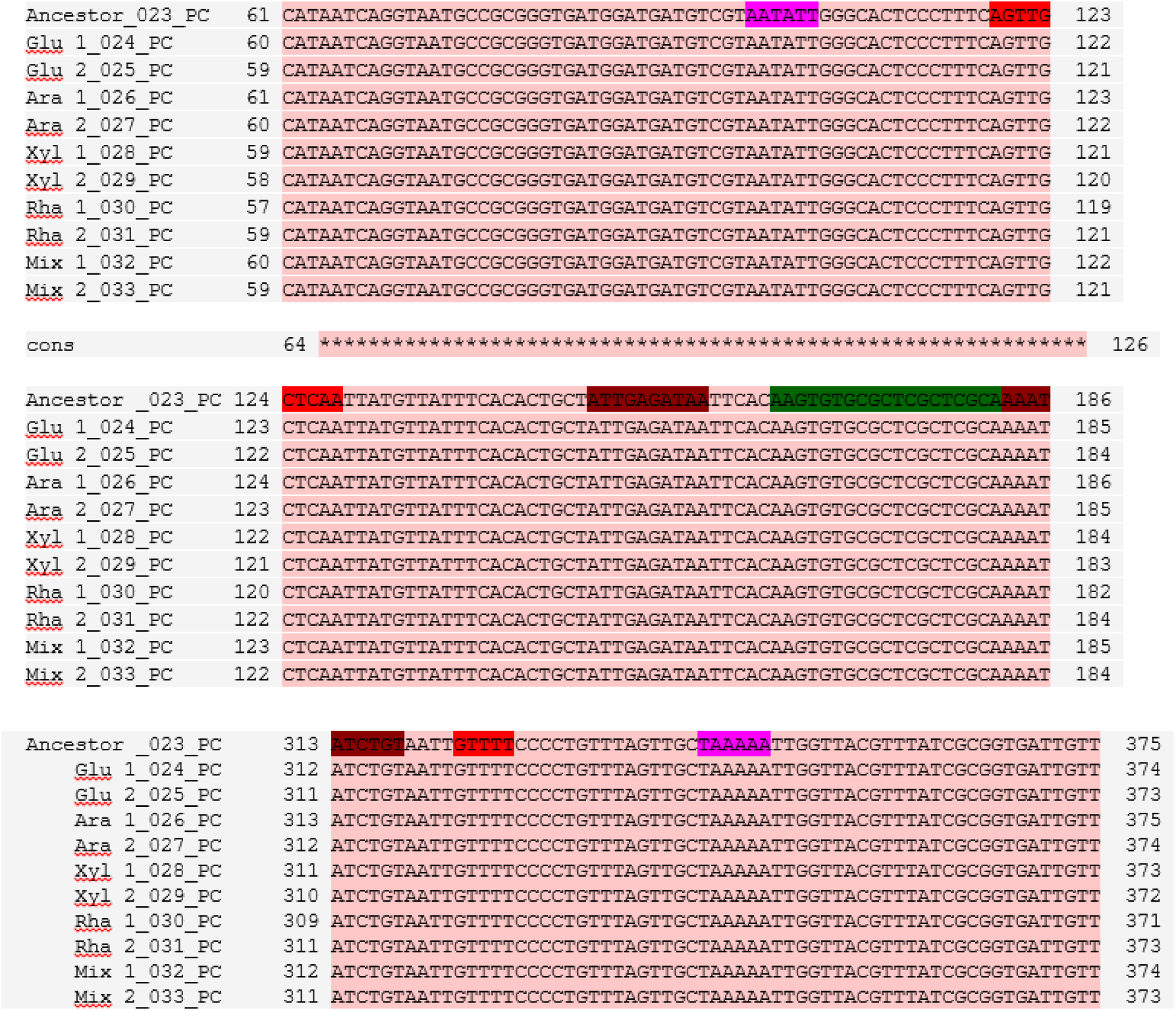
Sequence alignment data of *xylFGH* regulatory region in ancestor and evolved strains showing the XylR/AraC binding site -IA/IF (brown), CRP binding site (green) -10 (pink) and -35 (red) in (blue). *xylFGH* regulatory region was sequenced using cycle sequencing technology by Eurofins Bangalore and multiple sequence alignment was done in T-coffee alignment tool.

**Figure S10:**
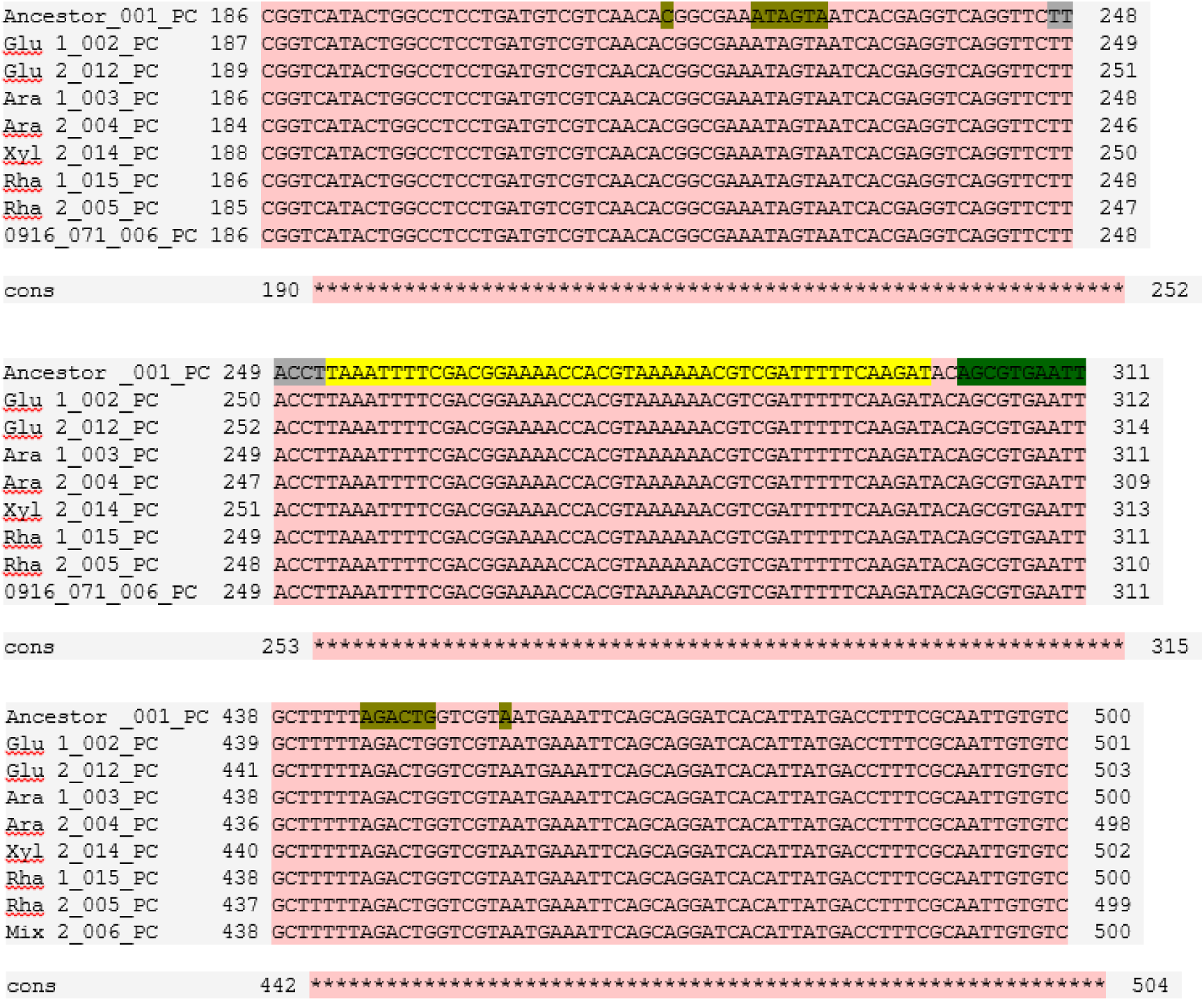
Sequence alignment data of *rhaBAD-rhaS* regulatory region in ancestor and evolved strains showing the RhaR binding site (yellow), RhaS binding site (pink) and CRP binding site (blue and brown). *rhaBAD-rhaS* regulatory region was sequenced using cycle sequencing technology by Eurofins Bangalore and multiple sequence alignment was done using T-coffee online tool.

**Figure S11:**
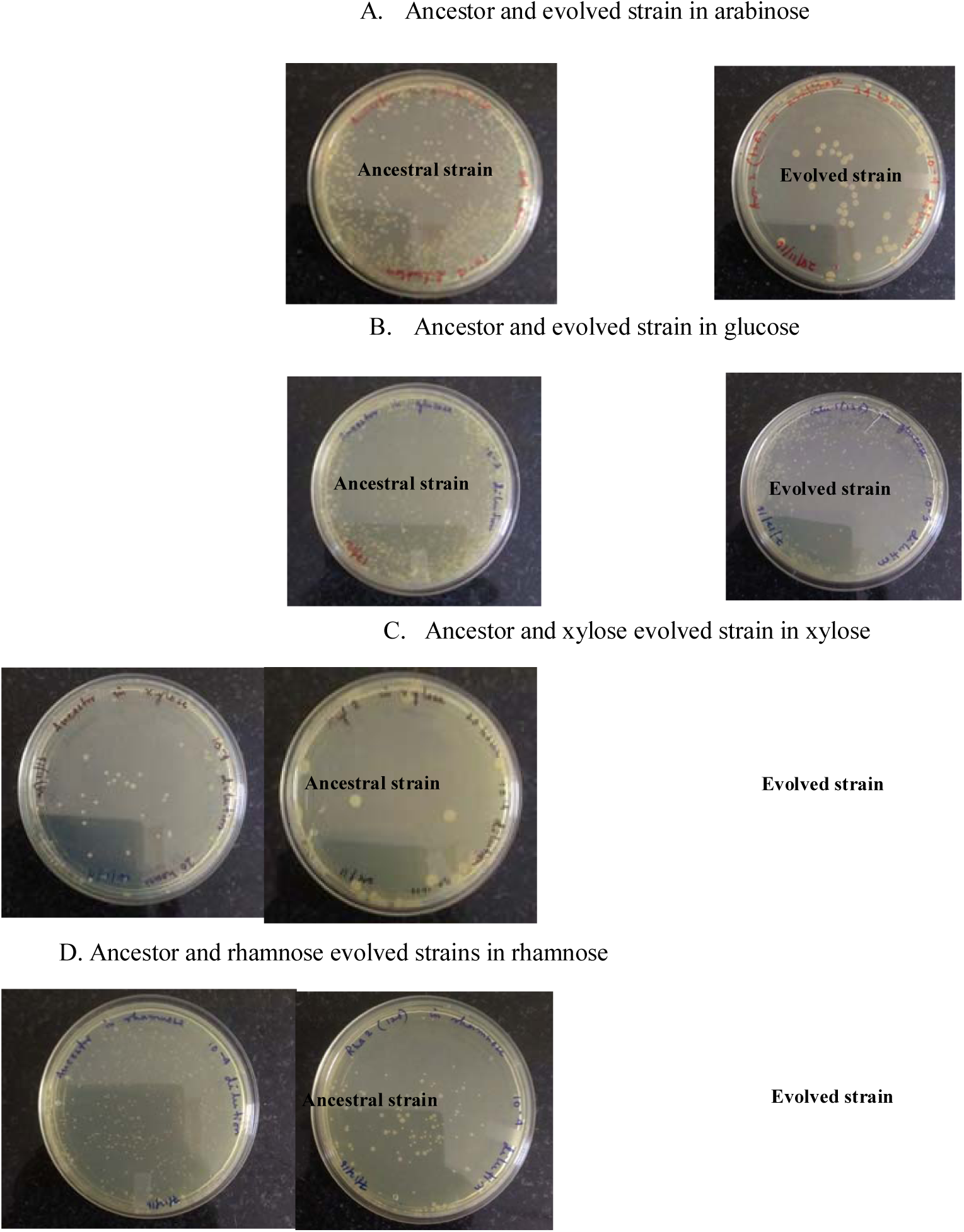
Growth of Ancestral and evolved strains in stationary phase. A. Ancestor and arabinose evolved strain B. Ancestor and glucose evolved strain C. Ancestor and xylose evolved strain D. Ancestor and rhamnose evolved strain. Ancestral and evolved strains were grow in adaptation environment till stationary phase. After which they were plated on LB agar and incubated till colonies appear.

**Table S1.**
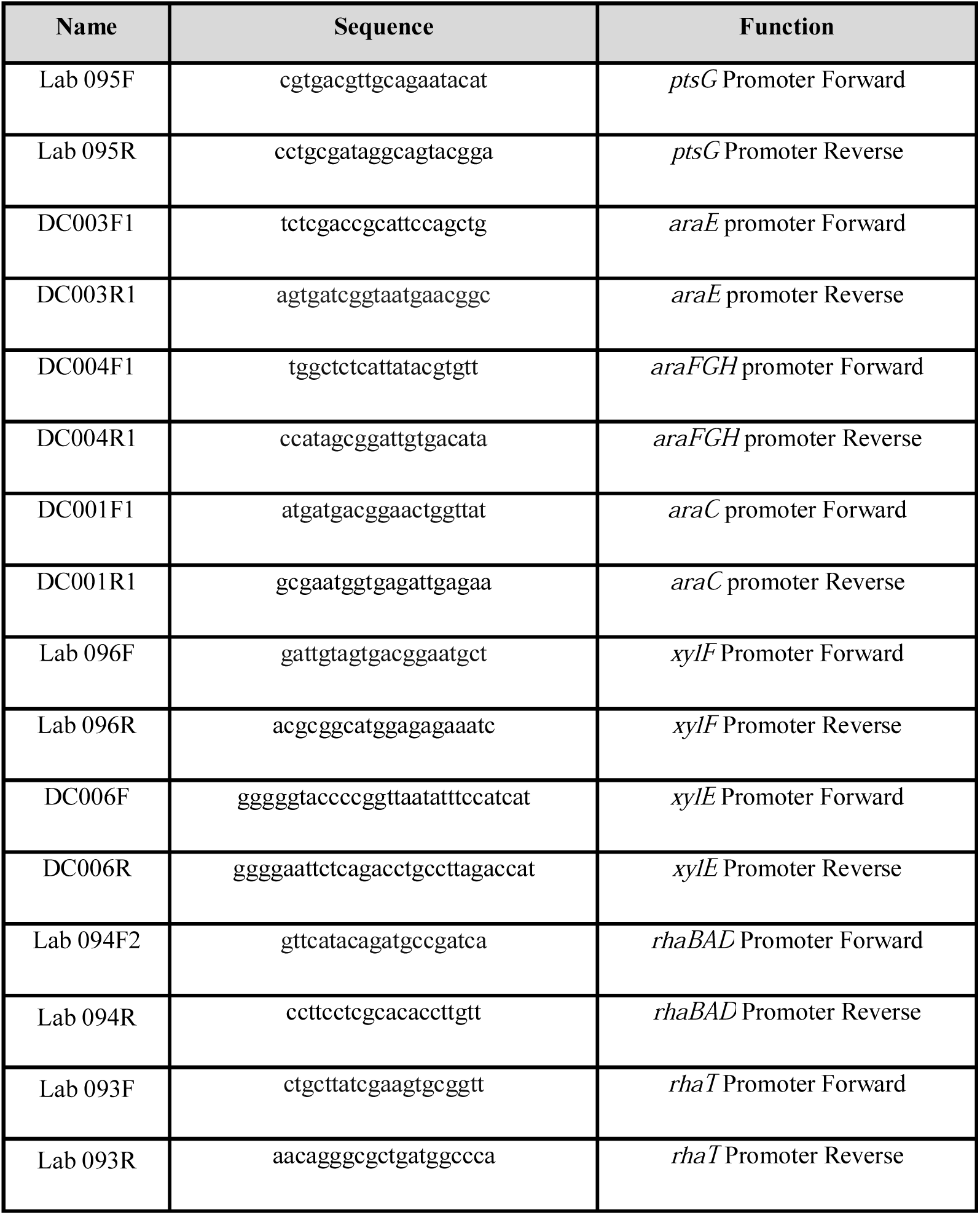
Primers used in this study

**Table S2A.** Lag phase duration, growth rate and steady state O.D. 600 values for ancestor and glucose evolved strains from different time points in non-selection environments

| Strain<br>(transfer<br>number) | Arabinose |  |  | Xylose |  |  | Rhamnose |  |  | Mixed- sugars |  |  |
| --- | --- | --- | --- | --- | --- | --- | --- | --- | --- | --- | --- | --- |
|  | Lag (h) | GR | OD | Lag (h) | GR | OD | Lag (h) | GR | OD | Lag (h) | GR | OD |
| **Ancestor** |  |  |  |  |  |  |  |  |  |  |  |  |
|  | 2 | 0.27 | 0.343 | 1.75 | 0.222 | 0.308 | 2.25 | 0.233 | 0.41 | 2.5 | 0.205 | 0.353 |
| **Glu 1** |  |  |  |  |  |  |  |  |  |  |  |  |
| (300) | 0.75<br>* | 0.281<br>ns | 0.329 | 0.75<br>* | 0.273<br>* | 0.331 | 0.75<br>** | 0.285<br>ns | 0.426 | 1.25<br>* | 0.309<br>* | 0.365 |
| (600) | 0.75<br>* | 0.29<br>ns | 0.338 | 0.75<br>* | 0.265<br>* | 0.322 | 0.75<br>** | 0.277<br>ns | 0.468 | 0.75<br>** | 0.275<br>** | 0.355 |
| (1200) | 0.75<br>* | 0.359<br>* | 0.331 | 0.75<br>* | 0.33<br>** | 0.314 | 0.75<br>** | 0.241<br>ns | 0.435 | 0.75<br>** | 0.337<br>** | 0.382 |
| (2000) | 0.75<br>* | 0.336<br>* | 0.336 | 0<br>** | 0.344<br>** | 0.325 | 0.75<br>** | 0.312<br>ns | 0.468 | 0.75<br>** | 0.353<br>** | 0.35 |
| **Glu 2** |  |  |  |  |  |  |  |  |  |  |  |  |
| (300) | 0.75<br>* | 0.283<br>ns | 0.332 | 0.75<br>* | 0.258<br>ns | 0.319 | 0.75<br>** | 0.281<br>ns | 0.407 | 1.125<br>* | 0.298<br>* | 0.342 |
| (600) | 0.75<br>* | 0.261<br>ns | 0.344 | 0.75<br>* | 0.282<br>* | 0.317 | 0.75<br>** | 0.26<br>ns | 0.441 | 0.75<br>** | 0.269<br>** | 0.345 |
| (1200) | 0.75<br>* | 0.277<br>* | 0.356 | 0.75<br>* | 0.193<br>** | 0.318 | 0.75<br>** | 0.192<br>ns | 0.394 | 0.75<br>** | 0.277<br>* | 0.356 |
| (2000) | 0.75<br>* | 0.336<br>** | 0.497 | 0<br>** | 0.368<br>** | 0.543 | 0.75<br>** | 0.278<br>ns | 0.46 | 0.75<br>** | 0.318<br>** | 0.373 |

**Table S2B.**
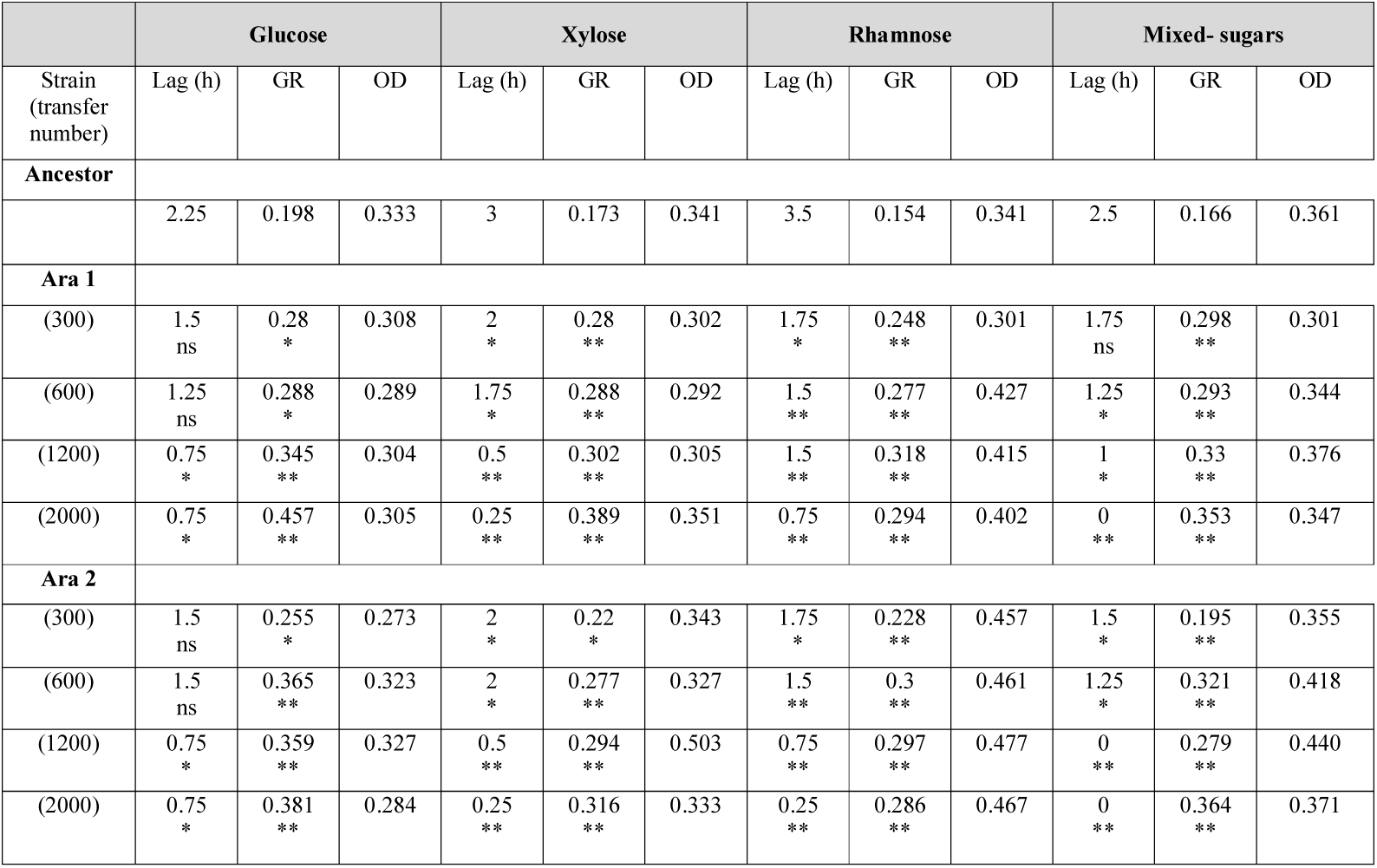
Lag phase duration, growth rate and steady state O.D. 600 values for ancestor and arabinose evolved strains from different time points in non-selection environments

**Table S2C.** Lag phase duration, growth rate and steady state O.D. 600 values for ancestor and xylose evolved strains from different time points in non-selection environments

|  | Glucose |  |  | Arabinose |  |  | Rhamnose |  |  | Mixed- sugars |  |  |
| --- | --- | --- | --- | --- | --- | --- | --- | --- | --- | --- | --- | --- |
| Strain<br>(transfer<br>number) | Lag<br>(h) | GR | OD | Lag (h) | GR | OD | Lag (h) | GR | OD | Lag (h) | GR | OD |
| **Ancestor** |  |  |  |  |  |  |  |  |  |  |  |  |
|  | 3.75 | 0.207 | 0.343 | 3.75 | 0.159 | 0.331 | 4.75 | 0.17 | 0.34 | 3.75 | 0.193 | 0.338 |
| **Xyl 1** |  |  |  |  |  |  |  |  |  |  |  |  |
| (300) | 3<br>* | 0.262<br>* | 0.298 | 3.25<br>ns | 0.243<br>** | 0.307 | 3<br>** | 0.249<br>** | 0.394 | 3<br>* | 0.286<br>* | 0.322 |
| (600) | 0.75<br>** | 0.289<br>** | 0.369 | 0.75<br>** | 0.288<br>** | 0.349 | 0.75<br>** | 0.248<br>** | 0.441 | 0.75<br>** | 0.305<br>** | 0.354 |
| (1200) | 0.75<br>** | 0.312<br>** | 0.315 | 0.75<br>** | 0.299<br>** | 0.357 | 0.75<br>** | 0.301<br>** | 0.435 | 0.75<br>** | 0.305<br>** | 0.364 |
| (2000) | 0.75<br>** | 0.327<br>** | 0.287 | 0.75<br>** | 0.228<br>** | 0.507 | 0.75<br>** | 0.21<br>ns | 0.486 | 0.75<br>** | 0.261<br>* | 0.348 |
| **Xyl 2** |  |  |  |  |  |  |  |  |  |  |  |  |
| (300) | 3.75<br>ns | 0.309<br>** | 0.313 | 3.75<br>ns | 0.259<br>** | 0.317 | 4<br>ns | 0.248<br>** | 0.391 | 4<br>ns | 0.29<br>* | 0.321 |
| (600) | 0.75<br>** | 0.308<br>** | 0.334 | 0.75<br>** | 0.311<br>** | 0.346 | 0.75<br>** | 0.237<br>** | 0.409 | 0.75<br>** | 0.304<br>** | 0.372 |
| (1200) | 0.75<br>** | 0.36<br>** | 0.319 | 0.75<br>** | 0.337<br>** | 0.336 | 0.75<br>** | 0.3<br>** | 0.435 | 0.75<br>** | 0.326<br>** | 0.376 |
| (2000) | 0.75<br>** | 0.363<br>** | 0.293 | 0.75<br>** | 0.212<br>** | 0.488 | 0.75<br>** | 0.218<br>ns | 0.487 | 0.75<br>** | 0.391<br>** | 0.357 |

**Table S2D.** Lag phase duration, growth rate and steady state O.D. 600 values for ancestor and rhamnose evolved strains from different time points in non-selection environments

|  | Glucose |  |  | Arabinose |  |  | Xylose |  |  | Mixed- sugars |  |  |
| --- | --- | --- | --- | --- | --- | --- | --- | --- | --- | --- | --- | --- |
| Strain<br>(transfer<br>number) | Lag (h) | GR | OD | Lag<br>(h) | GR | OD | Lag<br>(h) | GR | OD | Lag<br>(h) | GR | OD |
| **Ancestor** |  |  |  |  |  |  |  |  |  |  |  |  |
|  | 4.75 | 0.205 | 0.332 | 4.75 | 0.2 | 0.327 | 4.75 | 0.164 | 0.297 | 4.75 | 0.201 | 0.345 |
| **Rha 1** |  |  |  |  |  |  |  |  |  |  |  |  |
| (300) | 2.25<br>** | 0.339<br>** | 0.318 | 2.75<br>** | 0.284<br>* | 0.383 | 3<br>** | 0.279<br>* | 0.311 | 2.75<br>** | 0.258<br>** | 0.342 |
| (600) | 0.75<br>** | 0.373<br>** | 0.317 | 1<br>** | 0.313<br>** | 0.357 | 1.5<br>** | 0.264<br>ns | 0.285 | 0.75<br>** | 0.242<br>* | 0.302 |
| (1200) | 0.75<br>** | 0.348<br>** | 0.318 | 1<br>** | 0.338<br>** | 0.349 | 0.75<br>** | 0.314<br>** | 0.328 | 1<br>** | 0.254<br>* | 0.425 |
| (2000) | 0.75<br>** | 0.41<br>** | 0.313 | 1<br>** | 0.352<br>** | 0.331 | 0.75<br>** | 0.343<br>** | 0.338 | 0.75<br>** | 0.41<br>** | 0.360 |
| **Rha2** |  |  |  |  |  |  |  |  |  |  |  |  |
| (300) | 3<br>** | 0.326<br>** | 0.339 | 2.75<br>** | 0.271<br>* | 0.369 | 3.75<br>** | 0.364<br>** | 0.338 | 3 | 0.337<br>** | 0.358 |
| (600) | 0.75<br>** | 0.334<br>** | 0.308 | 1<br>** | 0.292<br>* | 0.339 | 0.75<br>** | 0.310<br>** | 0.331 | 0.75<br>** | 0.234<br>ns | 0.337 |
| (1200) | 0.75<br>** | 0.327<br>** | 0.332 | 0.75<br>** | 0.323<br>** | 0.431 | 0.75<br>** | 0.332<br>** | 0.324 | 0.75<br>** | 0.368<br>** | 0.399 |
| (2000) | 0.75<br>** | 0.373<br>** | 0.330 | 1.125<br>** | 0.372<br>** | 0.396 | 1.25<br>** | 0.345<br>** | 0.330 | 1.25<br>** | 0.352<br>** | 0.369 |

**Table S2E.** Lag phase duration, growth rate and steady state O.D. 600 values for ancestor and strains evolved in sugar-mixtures from different time points in non-selection environments

|  | Glucose |  |  | Arabinose |  |  | Xylose |  |  | Rhamnose |  |  |
| --- | --- | --- | --- | --- | --- | --- | --- | --- | --- | --- | --- | --- |
| Strain<br>(transfer<br>number) | Lag<br>(h) | GR | OD | Lag<br>(h) | GR | OD | Lag<br>(h) | GR | OD | Lag<br>(h) | GR | OD |
| **Ancestor** |  |  |  |  |  |  |  |  |  |  |  |  |
|  | 1.75 | 0.149 | 0.343 | 2 | 0.161 | 0.331 | 2 | 0.15 | 0.311 | 2 | 0.15 | 0.34 |
| **Mix1** |  |  |  |  |  |  |  |  |  |  |  |  |
| (300) | 1.5<br>ns | 0.274<br>** | 0.298 | 1.5<br>ns | 0.3<br>** | 0.307 | 1.125<br>ns | 0.291<br>** | 0.281 | 1.5<br>ns | 0.314<br>** | 0.394 |
| (600) | 1.25<br>ns | 0.24<br>** | 0.369 | 1.5<br>ns | 0.243<br>** | 0.349 | 1.5<br>ns | 0.265<br>** | 0.334 | 1.5<br>ns | 0.274<br>** | 0.441 |
| (1200) | 0.75<br>* | 0.385<br>** | 0.315 | 0.75<br>* | 0.273<br>* | 0.357 | 1<br>* | 0.293<br>** | 0.302 | 1.5<br>ns | 0.253<br>** | 0.435 |
| (2000) | 0.75<br>* | 0.364<br>** | 0.287 | 0.75<br>* | 0.328<br>** | 0.507 | 0.75<br>* | 0.332<br>** | 0.334 | 0.75<br>* | 0.316<br>** | 0.486 |
| **Mix2** |  |  |  |  |  |  |  |  |  |  |  |  |
| (300) | 2.25<br>ns | 0.273<br>** | 0.313 | 1.75<br>ns | 0.304<br>** | 0.317 | 1.125<br>ns | 0.275<br>** | 0.294 | 1.5<br>ns | 0.272<br>** | 0.391 |
| (600) | 1.5<br>ns | 0.252<br>** | 0.334 | 1.5<br>ns | 0.285<br>** | 0.346 | 1.875<br>ns | 0.281<br>** | 0.367 | 1.5<br>ns | 0.263<br>** | 0.409 |
| (1200) | 1.5<br>ns | 0.29<br>** | 0.319 | 1<br>* | 0.340<br>** | 0.336 | 1.5<br>ns | 0.314<br>** | 0.322 | 1.5<br>ns | 0.268<br>** | 0.435 |
| (2000) | 0.75<br>* | 0.364<br>** | 0.293 | 0.75<br>* | 0.389<br>** | 0.488 | 0.75<br>* | 0.373<br>** | 0.334 | 0.75<br>* | 0.331<br>** | 0.487 |

## References

Afroz, T., K. Biliouris, Y. Kaznessis and C. L. Beisel (2014). “Bacterial sugar utilization gives rise to distinct single-cell behaviors.” Mol Microbiol93(6): 1093–1103.

Aidelberg, G., B. D. Towbin, D. Rothschild, E. Dekel, A. Bren and U. Alon (2014). “Hierarchy of non-glucose sugars in Escherichia coli.” BMC Syst Biol8: 133.

Bachmann, H., M. Fischlechner, I. Rabbers, N. Barfa, F. Branco dos Santos, D. Molenaar and B. Teusink (2013). “Availability of public goods shapes the evolution of competing metabolic strategies.” Proc Natl Acad Sci U S A110(35): 14302–14307.

Baldoma, L. and J. Aguilar (1988). “Metabolism of L-fucose and L-rhamnose in Escherichia coli: aerobic-anaerobic regulation of L-lactaldehyde dissimilation.” J Bacteriol170(1): 416–421.

Beisel, C. L. and G. Storz (2011). “The base-pairing RNA spot 42 participates in a multioutput feedforward loop to help enact catabolite repression in Escherichia coli.” Mol Cell41(3): 286–297.

Bruckner, R. and F. Titgemeyer (2002). “Carbon catabolite repression in bacteria: choice of the carbon source and autoregulatory limitation of sugar utilization.” FEMS Microbiol Lett209(2): 141– 148.

Chen, Y. M., J. F. Tobin, Y. Zhu, R. F. Schleif and E. C. Lin (1987). “Cross-induction of the L-fucose system by L-rhamnose in Escherichia coli.” J Bacteriol169(8): 3712–3719.

David, J. D. and H. Weismeyer (1970). “Control of xylose metabolism in Escherichia coli.” Biochimica et Biophysica Acta (BBA) - General Subjects 201(3): 497–499.

Dekel, E. and U. Alon (2005). “Optimality and evolutionary tuning of the expression level of a protein.” Nature436(7050): 588–592.

Desai, T. A. and C. V. Rao (2010). “Regulation of Arabinose and Xylose Metabolism in Escherichia coli.” Applied and Environmental Microbiology 76(5): 1524–1532.

Egan, S. M. and R. F. Schleif (1993). “A Regulatory Cascade in the Induction of rhaBAD.” Journal of Molecular Biology 234(1): 87–98.

Gefen, O., O. Fridman, I. Ronin and N. Q. Balaban (2014). “Direct observation of single stationary-phase bacteria reveals a surprisingly long period of constant protein production activity.” Proceedings of the National Academy of Sciences 111(1): 556–561.

Gorke, B. and J. Stulke (2008). “Carbon catabolite repression in bacteria: many ways to make the most out of nutrients.” Nat Rev Microbiol 6(8): 613–624.

Haldimann, A. and B. L. Wanner (2001). “Conditional-replication, integration, excision, and retrieval plasmid-host systems for gene structure-function studies of bacteria.” J Bacteriol 183(21): 6384– 6393.

Hasona, A., Y. Kim, F. G. Healy, L. O. Ingram and K. T. Shanmugam (2004). “Pyruvate Formate Lyase and Acetate Kinase Are Essential for Anaerobic Growth of Escherichia coli on Xylose.” J Bacteriol 186(22): 7593-7600.

Heller, K. B., E. C. Lin and T. H. Wilson (1980). “Substrate specificity and transport properties of the glycerol facilitator of Escherichia coli.” J Bacteriol 144(1): 274–278.

Hernandez-Montalvo, V., F. Valle, F. Bolivar and G. Gosset (2001). “Characterization of sugar mixtures utilization by an Escherichia coli mutant devoid of the phosphotransferase system.” Appl Microbiol Biotechnol 57(1-2): 186–191.

Iida, A., S. Harayama, T. Iino and G. L. Hazelbauer (1984). “Molecular cloning and characterization of genes required for ribose transport and utilization in Escherichia coli K-12.” J Bacteriol 158(2): 674– 682.

Inada, T., K. Kimata and H. Aiba (1996). “Mechanism responsible for glucose-lactose diauxie in Escherichia coli: challenge to the cAMP model.” Genes Cells 1(3): 293–301.

Johnson, C. M. and R. F. Schleif (1995). “In vivo induction kinetics of the arabinose promoters in Escherichia coli.” J Bacteriol 177(12): 3438–3442.

Kassen, R. (2002). “The experimental evolution of specialists, generalists, and the maintenance of diversity.” Journal of Evolutionary Biology 15(2): 173–190.

Khankal, R., J. W. Chin and P. C. Cirino (2008). “Role of xylose transporters in xylitol production from engineered Escherichia coli.” J Biotechnol 134(3-4): 246–252.

Koirala, S., X. Wang and C. V. Rao (2015). “Reciprocal Regulation of l-Arabinose and d-Xylose Metabolism in Escherichia coli.” J Bacteriol 198(3): 386–393.

Kresge, N., R. D. Simoni and R. L. Hill (2005). “Otto Fritz Meyerhof and the elucidation of the glycolytic pathway.” J Biol Chem 280(4): e3.

Lawlis, V. B., M. S. Dennis, E. Y. Chen, D. H. Smith and D. J. Henner (1984). “Cloning and sequencing of the xylose isomerase and xylulose kinase genes of Escherichia coli.” Appl Environ Microbiol 47(1): 15–21.

Lenski, R. E., J. A. Mongold, P. D. Sniegowski, M. Travisano, F. Vasi, P. J. Gerrish and T. M. Schmidt (1998). “Evolution of competitive fitness in experimental populations of E. coli: what makes one genotype a better competitor than another?” Antonie Van Leeuwenhoek 73(1): 35–47.

Luo, Y., T. Zhang and H. Wu (2014). “The transport and mediation mechanisms of the common sugars in Escherichia coli.” Biotechnol Adv 32(5): 905–919.

Madar, D., E. Dekel, A. Bren, A. Zimmer, Z. Porat and U. Alon (2013). “Promoter activity dynamics in the lag phase of Escherichia coli.” BMC Syst Biol 7: 136.

Maddamsetti, R., P. J. Hatcher, A. G. Green, B. L. Williams, D. S. Marks and R. E. Lenski (2017). “Core Genes Evolve Rapidly in the Long-Term Evolution Experiment with Escherichia coli.” Genome Biol Evol 9(4): 1072–1083.

Makman, R. S. and E. W. Sutherland (1965). “ADENOSINE 3′,5′-PHOSPHATE IN ESCHERICHIA COLI.” J Biol Chem 240: 1309-1314.

Mayer, C. and W. Boos (2005). “Hexose/Pentose and Hexitol/Pentitol Metabolism.” EcoSal Plus 1(2).

Notredame, C., D. G. Higgins and J. Heringa (2000). “T-coffee: a novel method for fast and accurate multiple sequence alignment1.” Journal of Molecular Biology 302(1): 205–217.

Novak, M., T. Pfeiffer, R. E. Lenski, U. Sauer and S. Bonhoeffer (2006). “Experimental tests for an evolutionary trade-off between growth rate and yield in E. coli.” Am Nat 168(2): 242–251.

Okada, T., K. Ueyama, S. Niiya, H. Kanazawa, M. Futai and T. Tsuchiya (1981). “Role of inducer exclusion in preferential utilization of glucose over melibiose in diauxic growth of Escherichia coli.” Journal of Bacteriology 146(3): 1030–1037.

Oxman, E., U. Alon and E. Dekel (2008). “Defined order of evolutionary adaptations: experimental evidence.” Evolution 62(7): 1547–1554.

Schleif, R. (2000). “Regulation of the L-arabinose operon of Escherichia coli.” Trends Genet 16(12): 559–565.

Schneider, D., E. Duperchy, E. Coursange, R. E. Lenski and M. Blot (2000). “Long-term experimental evolution in Escherichia coli. IX. Characterization of insertion sequence-mediated mutations and rearrangements.” Genetics 156(2): 477–488.

Shimada, T., N. Fujita, K. Yamamoto and A. Ishihama (2011). “Novel Roles of cAMP Receptor Protein (CRP) in Regulation of Transport and Metabolism of Carbon Sources.” PLOS ONE 6(6): e20081.

Song, S. and C. Park (1997). “Organization and regulation of the D-xylose operons in Escherichia coli K-12: XylR acts as a transcriptional activator.” J Bacteriol 179(22): 7025–7032.

van Elsas, J. D., A. V. Semenov, R. Costa and J. T. Trevors (2011). “Survival of Escherichia coli in the environment: fundamental and public health aspects.” ISME J 5(2): 173–183.

Via, P., J. Badia, L. Baldoma, N. Obradors and J. Aguilar (1996). “Transcriptional regulation of the Escherichia coli rhaT gene.” Microbiology 142 (Pt 7): 1833–1840.

Winfield, M. D. and E. A. Groisman (2003). “Role of Nonhost Environments in the Lifestyles of Salmonella and Escherichia coli.” Applied and Environmental Microbiology 69(7): 3687–3694.

Woods, R., D. Schneider, C. L. Winkworth, M. A. Riley and R. E. Lenski (2006). “Tests of parallel molecular evolution in a long-term experiment with Escherichia coli.” Proc Natl Acad Sci U S A 103(24): 9107–9112.

Zwietering, M. H., F. M. Rombouts and K. van ’t Riet (1992). “Comparison of definitions of the lag phase and the exponential phase in bacterial growth.” J Appl Bacteriol 72(2): 139–145.

